# The actomyosin cortex controls t-tubule remodeling in muscle

**DOI:** 10.1101/2024.08.13.607563

**Authors:** Ana Raquel Pereira, Ana da Rosa Soares, Silvia Di Francescantonio, Tianyang Liu, Filomena A. Carvalho, Josie Liane Ferreira, Graciano Leal, Inês Faleiro, Naoko Kogata, Nuno C. Santos, Michael Way, Carolyn A. Moores, Edgar R. Gomes

## Abstract

Muscle cells form a network of plasma membrane invaginations called t-tubules that control calcium release from the endoplasmic reticulum at the triads during muscle contraction. Although the importance of t-tubules for muscle physiology is well established, and abnormalities are found in multiple disorders, the mechanisms that mediate t-tubule growth are unknown. We show that the actomyosin cortex beneath the plasma membrane, regulated by Arp2/3 complexes containing Arpc5, acts as a gatekeeper for the membrane availability required for t-tubule growth. Enlarged t-tubules are formed upon disruption of Arpc5, impairing the synchronisation between plasma membrane depolarization and calcium release. Additionally, ablation of Arpc5 postnatally in myofibers results in muscle fatigue and t-tubule abnormalities, as observed in muscle disorders. We propose that the actomyosin cortex impacts muscle function, offering a potential pathophysiological mechanism for muscle disorders.

## Main text

The contractile properties of skeletal muscle are essential for body movement, posture, and breathing. These vital functions are affected by multiple disorders and decline with age (*1*). Sarcomeres are the contractile units of muscle cells and contract synchronously upon calcium release from the endoplasmic reticulum (ER), also known as sarcoplasmic reticulum. Muscle contraction is initiated at the neuromuscular junction by local depolarisation of the plasma membrane, which propagates throughout the muscle cell (myofiber) surface (*2*). Since myofibers are the longest and thickest cells in the human body, the depolarisation signal is propagated throughout their entire volume via plasma membrane invaginations called transverse tubules (t-tubules) that surround sarcomeres (*3–5*). These t-tubules form membrane contact sites with two adjacent endoplasmic reticulum compartments, creating a structure termed a triad. Depolarisation of the t-tubule membrane triggers calcium release from the ER at the triads, a phenomenon termed excitation-contraction coupling (EC-coupling). Disruptions in triad assembly and morphology result in EC-coupling abnormalities, characteristic of multiple muscle disorders (*6*).

During mammalian development, a network of t-tubules emerges in myofibers in the embryo and becomes organized transversely only after birth (*7*). T-tubule biogenesis is initiated from plasma membrane invaginations mediated by Amphiphysin 2 (Bin1), a BAR domain protein, and caveolin-3-containing caveolae (*8*). Endosomal membrane traffic has been implicated in t-tubule network formation and maintenance (*3*, *9*). In adults, t-tubules have a surface area approximately seven times that of the myofiber plasma membrane, however it remains unclear how t-tubules grow (*10*).

The plasma membrane is tethered to the actomyosin cortex by membrane-to-cortex attachment proteins such as ezrin, radixin and moesin (ERM) family, that impact membrane remodelling (*11–13*). The actomyosin cortex consists of both linear and branched actin filaments, the latter of which is nucleated by the Arp2/3 complex composed of 7 evolutionarily conserved subunits (*14*, *15*). Notably, three Arp2/3 subunits are encoded by more than one gene in mammals, yielding Arp2/3 complexes with different properties (*16–18*). This diversity generates Arp2/3 complexes that appear to be tailored for specific cellular functions, including cell migration, viral transport, or T-cell activation (*16*, *19*, *20*). In skeletal muscle, we previously demonstrated that Arp2/3 complexes containing Arpc5L drive nuclei positioning to the periphery of myofiber by promoting desmin-dependent myofibril crosslinking to generate centripetal forces on the nucleus (*21*). In contrast, Arp2/3 complexes containing Arpc5 plays an essential role in triad organisation by an unknown mechanism. Here we investigated how Arpc5-containing Arp2/3 complexes are involved in t-tubule and triad organisation.

## Results

### Inhibition of the Arp2/3 complex promotes t-tubule growth

To understand how the t-tubule network is established in myofibers, we first evaluated t-tubule dynamics. For this, we incubated myofibers isolated from adult mice or *in vitro* developing myofibers with a plasma membrane dye (CellMask) to label t-tubules, and performed time-lapse microscopy (*7*). We observed that t-tubules exhibit growing and shrinking movements in both isolated myofibers and developing myofibers (Fig. 1A and 1B, Movie S1-S2). T-tubule growth was independent of t-tubule-ER contacts (fig. S1, Movie S3).

**Fig. 1.**
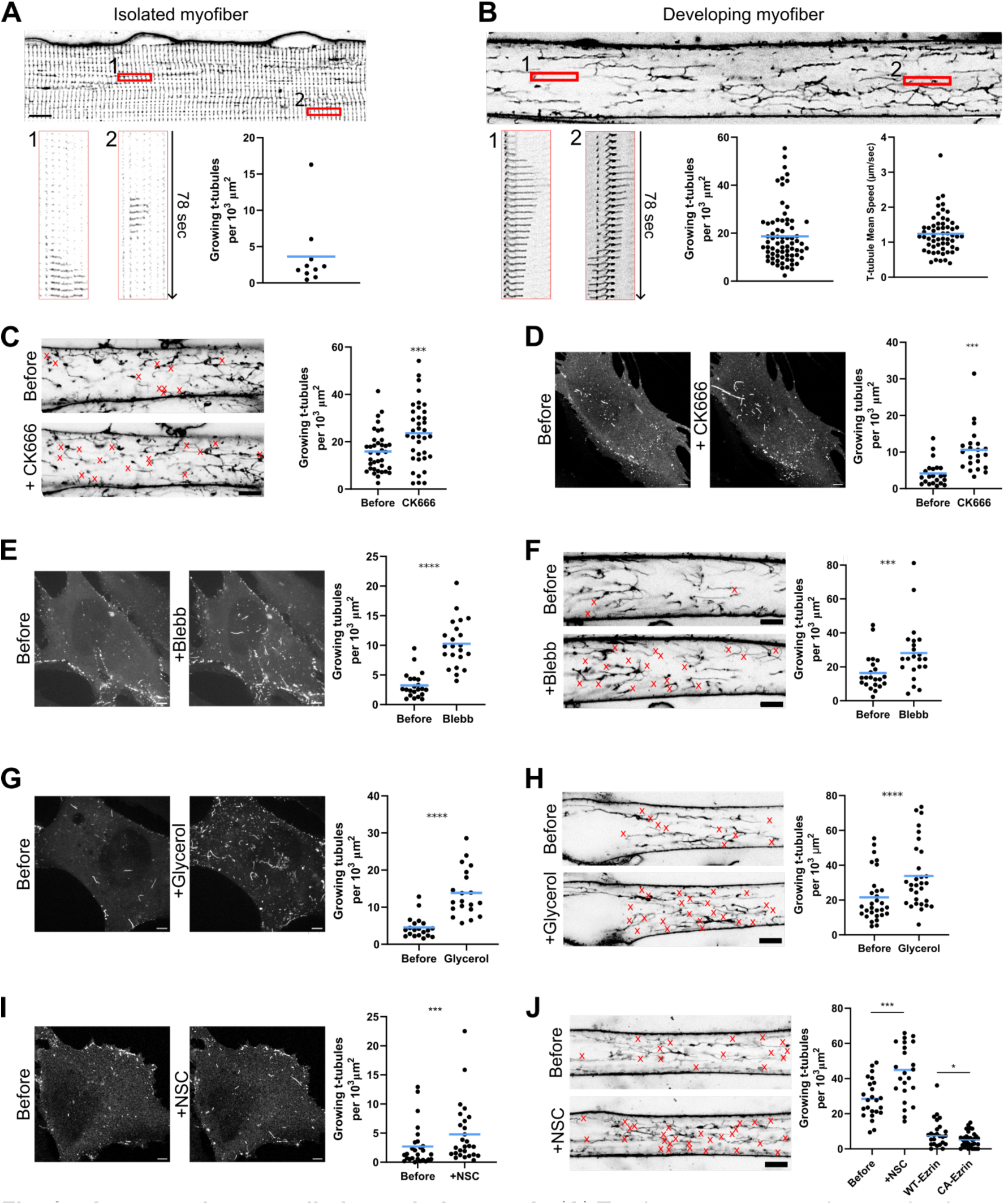
Actomyosin cortex limits t-tubule growth. (**A**) Top image corresponds to a timelapse snapshot of a mature isolated myofiber, stained with CellMask to label t-tubules and imaged by timelapse every 3 s for 2 min. Bottom image corresponds to kymographs from regions of interest (ROI) 1 and 2 of representative dynamic t-tubules. The plot represents the number of growing t-tubules per cell area during 2 min (N=10 isolated fibers, from three independent experiments). (**B**) Top image corresponds to a timelapse snapshot of an in vitro developing myofiber stained with CellMask to label t-tubules and imaged by timelapse every 3 sec for 2 min. At this stage of development (differentiation day 4), t-tubules are starting to form and have a longitudinal orientation. Bottom image corresponds to kymographs from ROIs 1 and 2 of representative dynamic t-tubules. The plots represent the number of growing t-tubules per cell area during 2 min (N=72 cells, from three independent experiments) and t-tubule growth mean speed (N=58 growing t-tubules, from three independent experiments). (**C**) Timelapse snapshots of in vitro developing myofibers (differentiation day 4) with CellMask labeled t-tubules before or after CK666 addition. Red X indicates the site of t-tubule growth during 2 min. The plot corresponds to the number of growing t-tubules per cell area during 2 min in the same myofibers before or after CK666 addition. Statistics: Wilcoxon test, p-value=0.0001; N=37 myofibers, results are from three independent experiments. (**D**) Timelapse snapshots of GFP-Bin1 before or after adding CK666. The plot corresponds to the number of growing tubules per cell during 2 min before or after adding CK666. Statistics: Wilcoxon test; N=22; results are from three independent experiments. (**E**) Timelapse snapshots of GFP-Bin1 myoblasts before or after para-aminoblebbistatin addition (Blebb). The plot corresponds to the number of growing tubules per cell area for 2 min before or after para-aminoblebbistatin addition. Statistics: Wilcoxon test; N=22 cells, results are from three independent experiments. (**F**) Timelapse snapshots of in vitro developing myofibers (differentiation day 4) with CellMask labeled t-tubules before or after para-aminoblebbistatin addition (Blebb). Red X indicates the site of t-tubule growth during 2 min. The plot corresponds to the number of growing t-tubules per cell area during 2 min in the same myofibers before or after para-aminoblebbistatin addition. Statistics: Wilcoxon test, p-value=0.0001; N=22 myofibers, results are from three independent experiments. (**G**) Timelapse snapshots of GFP-Bin1 myoblasts before or after glycerol addition. The plot corresponds to the number of growing tubules per cell area for 2 min before or after glycerol addition. Statistics: Wilcoxon test; N=20 cells, results are from three independent experiments. (**H**) Timelapse snapshots of in vitro developing myofibers (differentiation day 4) with CellMask labeled t-tubules before or after glycerol addition. Red X indicates the site of t-tubule growth during 2 min. The plot corresponds to the number of growing t-tubules per cell area during 2 min in the same myofibers before or after glycerol addition. Statistics: Wilcoxon test; N=30 myofibers, results are from three independent experiments. Scale bars 5 µm. (**I**) Timelapse snapshots of GFP-Bin1 myoblasts before or after NSC668394 (NSC). The plot corresponds to the number of growing tubules per cell area for 2 min before or after NSC addition. Statistics: Wilcoxon test; N=27 cells, results are from four independent experiments. (**J**) Timelapse snapshots of in vitro developing myofibers (differentiation day 4) with CellMask labeled t-tubules before or after NSC668394 addition (NSC). Red X indicates the site of t-tubule growth during 2 min. The plot corresponds to the number of growing t-tubules per cell area during 2 min in the same myofibers before NSC, after NSC, in cells expressing Ezrin-GFP (WT-Ezrin) or expressing constitutively active Ezrin-GFP (CA-Ezrin, T567D-GFP). Statistics: Before and After NSC: Paired t-test; N=24 myofibers. WT-Ezrin and CA-Ezrin: Mann-Whitney. WT-Ezrin N=25 and CA-Ezrin=33 myofibers. Results are from three independent experiments. Blue lines on all plots represent mean values. A-D, F-K scale bars 5 µm.

We previously found that triad organisation during myofiber formation depends on the Arp2/3 complex (*21*). Building upon this observation, we hypothesised that the Arp2/3 complex is required for t-tubule growth. We treated developing myofibers with the Arp2/3 inhibitor CK666, and contrary to our hypothesis, there was a significant increase in the number of growing t-tubules (Fig. 1C, Movie S4). T-tubule formation is mediated by Bin1 and its overexpression is sufficient to induce plasma membrane tubulation (*8*, *22*, *23*). Stable expression of GFP-Bin1 in C2C12 myoblasts (GFP-Bin1 myoblasts) resulted in dynamic tubules similar to those in developing myofibers (Movie S5). Moreover, treatment with CK666 also increased the number of growing tubules (Fig. 1D; Movie S5). Together, these results show that Arp2/3 inhibition promotes t-tubule and membrane tubule growth.

### F-Actin does not accumulate near growing tubules

Actin cytoskeleton and Arp2/3 complex can modulate membrane tubulation in multiple organelles (*24*). To investigate whether actin was enriched at growing tubules we imaged actin filaments (F-actin) using LifeAct-mCherry or mScarlet-Utrophin on membrane tubules formed in GFP-Bin1 myoblasts. We did not observe F-actin enrichment during tubule growth (fig. S2 A-D; Movie S6 and S7). We also performed cryo-correlative Light and Electron Microscopy (cryo-CLEM) and cryo-electron tomography (cryo-ET) of GFP-Bin1 myoblasts in regions containing tubules (fig. S2E-I, S3, S4). We observed unbranched F-actin forming bundles or meshes throughout the different volumes (fig. S2H, I, S4A and S4B, Movie S8-S10). However, analysis of the volume across different z positions revealed no distinct arrangement of actin filaments near the tubules when compared to other cytoplasmic regions (fig. S2I, S4C-J). Importantly, we observed very few actin branches around tubules (fig. S2I), and they were not more abundant when compared to other regions of the cytoplasm (fig. S4C-J). This stands in contrast to clathrin-mediated endocytosis, where branched actin is enriched near the endocytic membrane (*25*, *26*). Overall, these data suggest that branched F-actin does not accumulate near growing t-tubules, leading us to explore contributions by the Arp2/3 network to t-tubule dynamics elsewhere in the cell.

### The actomyosin cortex limits t-tubule growth

During myofiber development, plasma membrane must be available to form t-tubules. Plasma membrane is regulated by the underlying cell cortex consisting of an actomyosin network linked to the plasma membrane by the ezrin-radixin-moesin proteins (ERM) (*11*, *13*). Since the cortical actin is regulated by the Arp2/3 complex, we hypothesised that perturbing the actomyosin cortex leads to increased membrane availability required for t-tubule growth (*15*, *27*, *28*). To test this hypothesis, we interfered with actomyosin cortex by inhibiting myosin-2, using para-aminoblebbistatin (blebbistatin), and quantified the number of growing tubules before and after treatment (*29*, *30*). Blebbistatin treatment of GFP-Bin1 myoblasts or developing myofibers led to a significant increase in the number of growing tubules (Fig. 1E and 1F; Movie S11 and S12). We also induced an excess of plasma membrane by a rapid reduction of cell volume with a hyperosmotic shock (*31*). Quantification of the number of growing tubules before and after osmotic shock revealed an increase in the number of growing tubules (Fig. 1G and 1H, Movie S13 and S14). In addition, the interference of membrane to cortex attachment with NSC668394, an ezrin phosphorylation inhibitor (*32*) also results in an increase in the number of growing tubules (Fig. 1I and 1J; Movie S15 and S16). Conversely, overexpression of a constitutively active form of Ezrin (CA-Ezrin) (*33*) reduces the number of growing t-tubules (Fig. 1J, graph). Importantly, cells expressing CA-Ezrin had fewer growing t-tubules when compared to those with WT-Ezrin (Fig. 1J, graph). Together, these findings suggest that the actomyosin cortex functions as a regulatory gatekeeper in the growth of t-tubules.

### Arpc5 limits t-tubule growth through the actomyosin cortex

We showed that Arp2/3 complexes containing Arpc5 are involved in triad organization (*15*, *21*, *27*). Therefore, we hypothesized that Arpc5 role in triad organization relates to its function in regulating cortical actin-mediated t-tubule growth. To test if Arpc5 had a role in t-tubule growth, we performed knock-down (KD) experiments with siRNA targeting Arpc5 in GFP-Bin1 myoblasts and developing myofibers (fig. S5). We found that depletion of Arpc5, results in significantly more growing tubules in GFP-Bin1 myoblasts and t-tubules in developing myofibers when compared to controls (Fig. 2A and 2B; Movie S17 and S18). Next we evaluated the role of Arpc5 on the actomyosin cortex in developing myofibers. We found that non-muscle-myosin2B (NM-Myo2B) localizes as foci at the membrane that were reduced upon Arpc5 depletion (Fig. 2C and 2D). In addition, the levels of actin ERM proteins (pERM) were also reduced and its distribution was more heterogeneous following Arpc5 KD (Fig. 2E-G). Finally we assessed muscle cells’ nanoscale topology using Atomic Force Microscopy (AFM) and found that depletion of Arpc5 leads to an increase in membrane roughness (Fig. 2H). These data suggests that depletion of Arpc5 impacts the actomyosin cortex and the membrane to cortex attachment in developing myofibers (Fig. 2I).

**Fig. 2.**
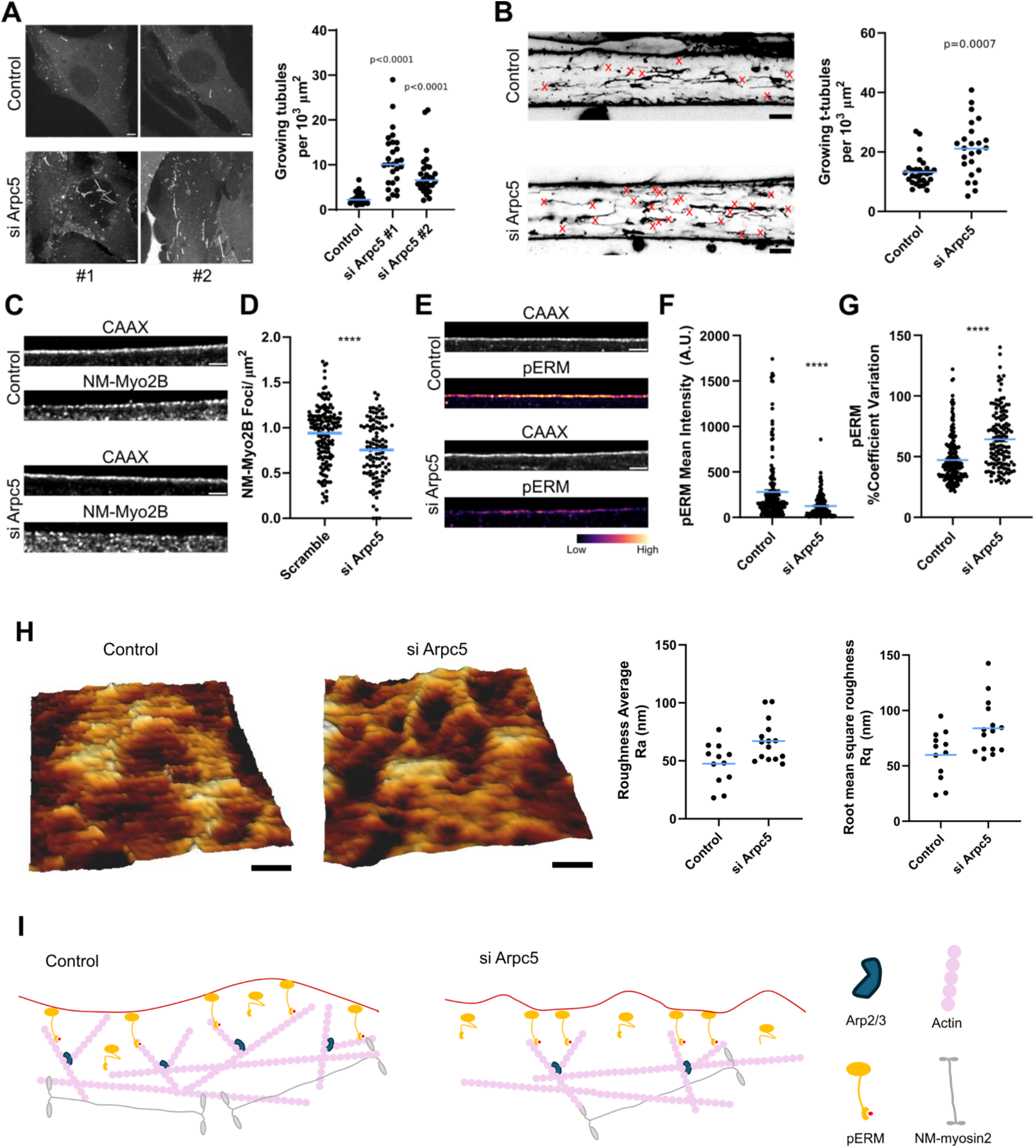
Arpc5 regulates cortical actin to limit t-tubule growth. **(A)** Timelapse snapshots of GFP-Bin1 myoblasts transfected with scramble (Control) or Arpc5 (si Arpc5) siRNA. The plot corresponds to the number of growing tubules per cell area during 2 min. Statistics: Mann-Whitney; Control N=25; siArpC5#1 N=27 siArpC5#2 N=28. Results from three independent experiments. Scale bars 5 µm. (**B**) Timelapse snapshots of in vitro developing myofibers (differentiation day 4) with CellMask-labelled t-tubules from scramble (Control) or Arpc5 depleted (siArp5) myofibers. Red X indicates sites of t-tubule growth during 2 min. The plot corresponds to the number of growing t-tubules per cell area during 2 min. Statistics: Mann-Whitney, p-value=0.0007; Control N=29; siArpc5 N=24; results are from three independent experiments. Scale bars 5 µm. (**C**) Non-muscle myosin 2B (NM-Myo2B) staining in CAAX-GFP myofibers from scramble (Control) or depleted for Arpc5 (si Arpc5). Scale bar 2 µm. (**D**) Quantification of NM-myo2B foci per ROI area (µm^2^). Each spot in the graph is the average number of identified NM-myo2B peaks per ROI segment area. Control N=164 ROIs, siArpc5 N=106. Statistics unpaired t-test, p-value <0.0001. Results are from three independent experiments. (**E**) PhosphoERM (pERM) staining in CAAX-GFP myofibers from scramble (Control) or depleted for Arpc5 (siArpc5). Scale bar 2 µm. (**F**) Plot corresponding to phosphoERM (pERM) mean intensity quantified over consecutive 10 µm length segments at the plasma membrane (each spot is the average of the pixel fluorescence intensity over the 10 µm segments). Control N=188 ROIs, siArpc5 N=148. Statistics Mann-Whitney, p-value <0.0001. Results are from three independent experiments. (**G**) Plot corresponding to the percentage of the coefficient of variation of phosphoERM signal intensity (pERM) over consecutive 10 µm segments at the plasma membrane. Control N=188 ROIs, siArpc5 N=148. Statistics Mann-Whitney, p-value <0.0001 (**H**) Atomic Force Microscopy (AFM) of developing myofibers. On the left a representative QI-mode height map image is shown for control and si Arpc5 depleted myofibers. Scale bar 0.2 µm. Height scale 200 nm. Graphs show the average roughness (Ra) and root mean square roughness (Rq) quantifications for surface topography. Surface topographies of control cells (N=12) and Arpc5 KD cells (N=15) were analyzed. Statistics Ra, unpaired t-test, p=0.0087. Statistics Rq, Student’s unpaired t-test, p=0.0143. Results are from three independent experiments (**I**) Schematic representation showing that Arpc5 is important for maintaining the membrane to cortex attachment and membrane flatness, gatekeeping membrane availability for t-tubule growth. Blue lines on all plots represent mean values.

### Arpc5 depletion leads to enlarged t-tubule clusters and asynchronous EC-coupling

During myofiber development, t-tubules form two-sided connections with the ER, establishing triads that become transversely organised (*3–5*). Since loss of Arpc5 impacted triad organisation (*21*), we investigated the role of Arpc5 in the t-tubule network in more differentiated myofibers (*21*, *34*). In control conditions, we observed transversely and longitudinally aligned t-tubules which are evenly distributed throughout the myofiber (Fig. 3A). In contrast, in Arpc5 KD myofibers, t-tubules formed clusters (Fig. 3A-C). Using joint-deconvolution Airyscan microscopy, we found that the clusters are composed of enlarged t-tubules (Fig. 3D). Transmission electron microscopy (TEM) of differentiated myofibers confirmed a higher proportion of enlarged t-tubules in Arpc5 KD myofibers (Fig. 3E and 3F). Furthermore, we observed a reduction of triads in Arpc5 KD compared to the control both using TEM and fluorescence microscopy of Junctophilin-1 (JPH1) puncta, a marker of triads, (Fig. 3E green arrowheads, 3G-I). Together, these results show that Arpc5 depletion impacts t-tubule and triad morphology.

**Fig. 3.**
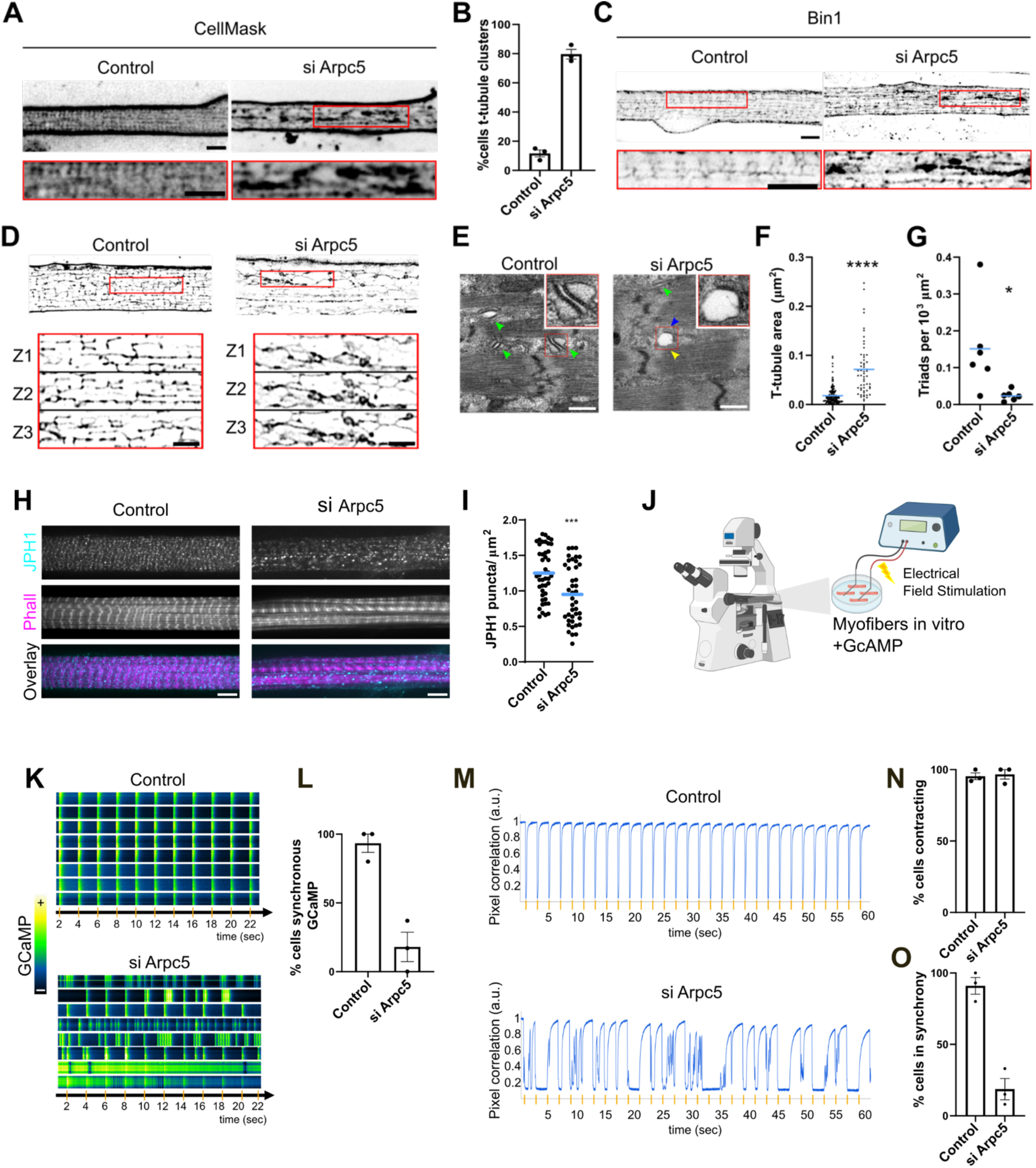
Arpc5 is involved in triad organization and synchronous EC-coupling. **(A)** Representative images of CellMask labelled t-tubules from siRNA treated control (scramble) or Arpc5 (si Arpc5) myofibers (differentiation day 7). Magnifications corresponding to the red boxes are shown below each image. Scale bars 5 μm. (**B**) Quantification of the percentage of myofibers with t-tubule clusters. Control N=42 myofibers; si Arpc5 N= 41 myofibers, from three independent experiments. **(C)** Representative images of Bin1 immunostained t-tubules from siRNA treated control (scramble) or Arpc5 (si Arpc5) depleted myofibers (differentiation day 7). Magnifications corresponding to the red boxes are shown below each image. Scale bars 5 μm (**D**) Higher resolution reconstructions of Airyscan images using a Joint Deconvolution method. Magnification Z slices corresponding to the red boxes are shown below of each image. Scale bars 2 μm. (**E**) Transmission Electron Microscopy (TEM) of scramble (Control) or Arpc5 (si Arpc5) depleted myofibers (differentiation day 7). Green arrowheads indicate triads (one t-tubule in contact with two adjacent ER compartments). Blue arrows indicate t-tubules connected to the ER only on one side, not forming a triad. Yellow arrowheads indicate enlarged t-tubules. Scale bars 500 nm. Magnifications corresponding to the red ROIs are shown as insets, scale bars 100 nm. (**F**) Plot representing the area of each t-tubule. Control N=162 t-tubules, si Arpc5 N=56 t-tubules. A total of 6 cells were quantified per condition, two biological replicates. Mann-Whitney test (p<0.0001). (**G**) Quantifications of the number of triads normalized for cell area (10^3^ μm^2^). Student’s t-test (p=0.0286). (**H**) Representative images of control (scramble), Arpc5 (si Arpc5) depleted myofibers (differentiation day 7) immunostained for Junctophilin 1 (JPH1, cyan), Bin1 (Yellow); and stained with phalloidin (Phall, magenta). Scale bars 5 µm. (**I**) Quantification of the number of Junctophilin 1 (JPH1) puncta per µm^2^ in scramble (Control) or Arpc5 (si Arpc5) depleted myofibers. Statistics: Mann-Whitney statistical test, siArpc5 p=0.0005; Control N=44 myofibers; si Arpc5 N=41 myofibers. Results are from three independent experiments. (**J**) To assess the impact of Arpc5 in EC-coupling, in vitro myofibers differentiated for 7-8 days expressing GCaMP are stimulated with 10 V at a frequency of 0.5 Hz to elicit membrane depolarization which ultimately results in calcium release and cell contraction. (**K**) Subcellular GCaMP fluorescence kymographs from 8 different myofibers of control (Scramble) or Arpc5 (siArpc5) are shown. Orange marks indicate electrical pulses. (**L**) Quantification of the percentage of myofibers from **K** with GCaMP fluorescence transients synchronised with the electrical pulse. Control N=40, si Arpc5 N=62. Results are from three independent experiments. (**M**) Representative contraction tracing recordings from control (Scramble) or Arpc5 (si Arpc5) depleted myofiber. Orange marks indicate electrical pulses. (**N**) Quantification of the percentage of contracting myofibers from **M**. Control N=43, siArpc5 N=36. Results are from three independent experiments. (**O**) Quantification of the percentage of myofibers from **M** that contract in synchrony with the electrical impulse. Control N=41, siArpc5 N=35. Results are from three independent experiments. Blue lines on all plots represent mean values. Black lines in all plots represent the standard error of the mean (SEM).

As triads are fundamental for EC-coupling, we assessed the impact of the loss of Arpc5 on calcium release and cell contraction in differentiated myofibers (Fig. 3J). We found that myofibers depleted for Arpc5, exhibited abnormalities in calcium transients, which were not synchronised with the electrical stimulation, when compared to WT myofibers (Fig. 3K and 3L). We then performed single-myofiber contraction tracing (Fig. 3M) and found that myofibers could still contract when Arpc5 is depleted (Fig. 3M and 3N). However, there was a dramatic asynchrony between electrical stimulation and myofiber contraction (Fig. 3M and 3O). These results highlight the role of Arpc5 containing Arp2/3 complexes and limiting t-tubule overgrowth in EC-coupling for muscle contraction.

### Arpc5 is associated muscle function during postnatal development in mice

Our in vitro myofibers are differentiated up to a stage equivalent to postnatal day 1 myofibers, which have nuclei at the periphery, organized transverse triads, and a network of transverse and longitudinal t-tubules (*35*, *36*). However, the t-tubule transverse network only becomes fully established during the first weeks of postnatal development in mice (*7*, *36*). To determine if Arpc5 has a role to further mature t-tubules and triads, we investigated mouse skeletal muscle postnatal development. As constitutive Arpc5 knockout mouse model is embryonically lethal (*37*), we used a tamoxifen-inducible, skeletal muscle-specific Arpc5 knockout model (Arpc5^-/-^, Fig. 4A) (*38*). To induce Arpc5 knockout during postnatal development, we administered 4OH-tamoxifen immediately after birth - postnatal (P) days 1, 2 and 3 - and characterized the neonatal Arpc5^-/-^ mice at P7. Arpc5 protein levels were significantly decreased when compared to Arpc5^fl/fl^ (Arpc5^WT^) (Fig. 4B and 4C). We performed TEM on longitudinal sections of hind limb muscle and found that the plasma membrane of Arpc5^-/-^ mice was more irregular than Arpc5^WT^ (Fig. 4D). Mature transverse t-tubules were decreased, whereas diagonal and longitudinal t-tubules did not change, suggesting that Arpc5 is also involved in t-tubule maturation that occurs postnatally. We also observed an increase of enlarged t-tubules in triads (Fig. 4E-G). Additionally, we found no difference in the number of triads between Arpc5^WT^ and Arpc5^-/-^ mice (Fig. 4H). Finally, structural alterations described above were associated with functional impairments in Arpc5^-/-^ skeletal muscle physiology, as hind limb suspension tests revealed increased fatigue and muscle weakness in Arpc5^-/-^ mice (Fig. 4I). These findings unveil the role of Arpc5 in t-tubule maturation and muscle function during postnatal development.

**Fig. 4.**
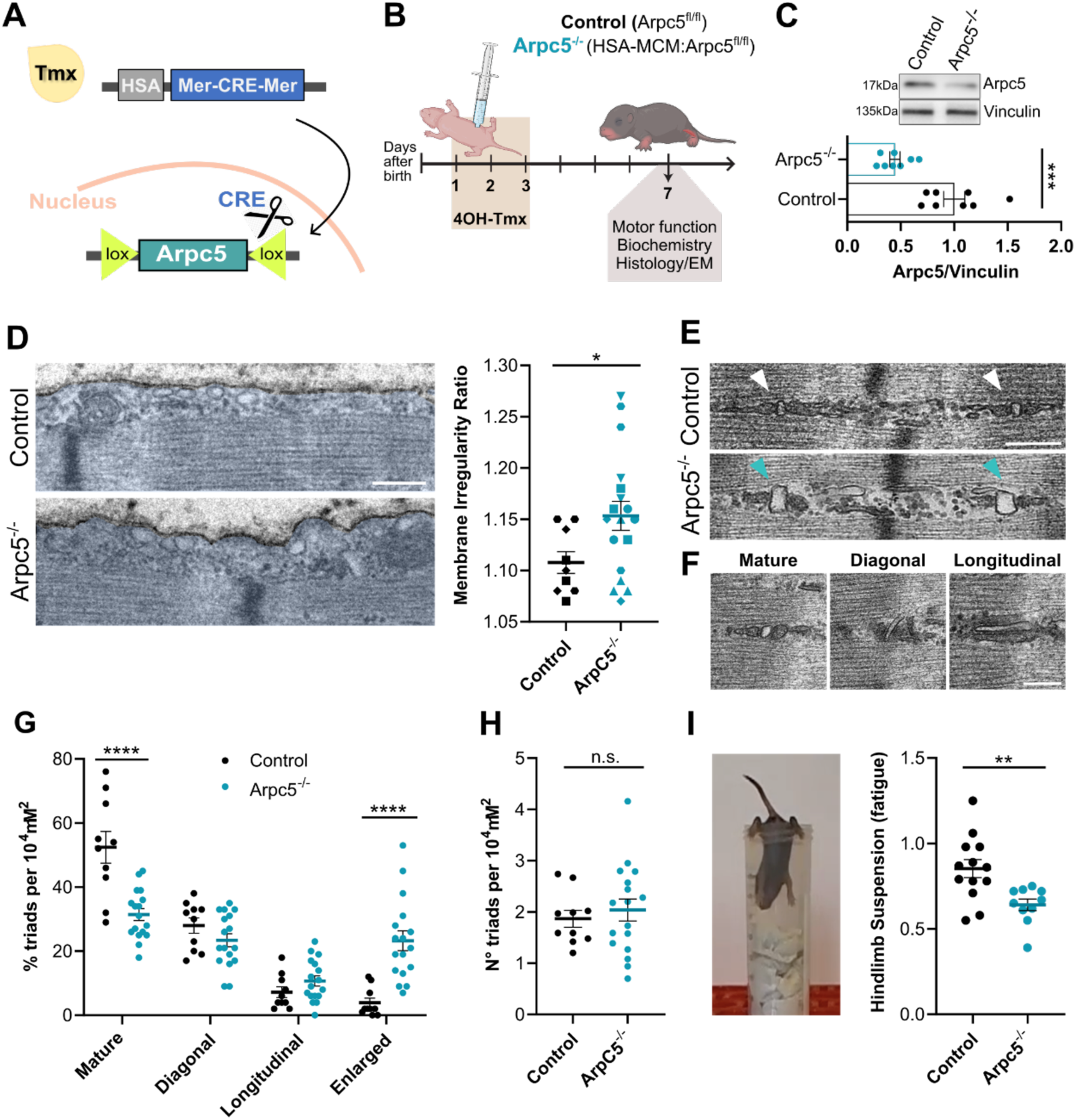
Postnatal depletion of Arpc5 compromises triad structure and muscle function. (**A**) Schematic of the Cre-lox knockout strategy to induce skeletal-muscle specific depletion of Arpc5. (**B**) Schematic of the inducible skeletal-muscle specific HSA-MCM mouse model used in this study. Daily 4OH-tamoxifen administration from P1 to P3 was used to induce skeletal muscle-specific Arpc5 depletion. At P7, locomotion was evaluated and the hindlimb posterior muscles were collected for TEM analysis. (**C**) Upper panel, representative Western blots showing Arpc5 and Vinculin levels in skeletal muscle of P7 Control and Arpc5^-/-^ mice. Lower panel, quantification of Western blots, showing a significant decrease of Arpc5 protein levels in Arpc5^-/-^ mice. The band intensity of each protein was normalized against vinculin, and the results are shown as fold vs the average of littermate controls. Statistic: unpaired 2-tailed Student’s t test, with Mann-Whitney test, p=0.0002. (**D**) Left panel, TEM representative images of longitudinal hindlimb muscle from control (Control) or Arpc5^-/-^ mice. The myofiber cytoplasm is pseudocolored in light blue. Note the darker contour line outlining the myofiber plasma membrane. Scale bar 500 nm. Right panel, quantification of membrane irregularity ratio in myofibers from P7 Control and Arpc5^-/-^ mice, measuring the deviation of a surface from a straight line. Values higher than 1 indicate a rougher membrane. Statistic: unpaired 2-tailed Student’s t test, with Welch’s correction, p=0.0157. (**E**) TEM representative images of myofibers from control (Control) or Arpc5^-/-^ mice. White arrowheads indicate mature triads (one transverse elongated t-tubule in contact with two adjacent ER compartments). Green arrows indicate enlarged t-tubules, forming an abnormal triad. Scale bar 250 nm. (**F**) Representative images of triads with t-tubule and ER with various morphologies - longitudinal, diagonal, and mature - indicated by white arrow and reflecting progressive changes in postnatal triad maturation. Scale bar 200 nm. (**G**) Quantifications of the percentage of diagonal, longitudinal, mature and enlarged triads normalized for cell area (10^4^ μm^2^) in myofibers from P7 Control and Arpc5^-/-^ mice. Statistic: 2-way ANOVA, Sidack’s multiple comparisons test; adjusted p value mature normal/enlarged triads <0.0001. (**H**) Quantifications of the number of all mature triads normalized for cell area (10^4^ μm^2^). Statistic: unpaired 2-tailed Student’s t test, non-parametric Mann–Whitney test was used, p>0.05. (**I**) Quantification of the fatigue shown by P7 Control and Arpc5^-/-^ neonates while performing the hindlimb suspension test. Fatigue was assessed by calculating the ratio between the time to fall in the final and initial measurements during the hindlimb suspension test. A lower fatigue value indicates greater fatigue in the mouse. Control N=13 Arpc5^-/-^ N=10 mice. Statistic: unpaired 2-tailed Student’s t test, with Welch’s correction, p=0.0032. Black horizontal lines in all plots represent mean values. Black vertical lines in all graphs represent SEM.

## Conclusions

T-tubules and their connection with the endoplasmic reticulum to form triads are fundamental for muscle contraction and are disrupted in multiple muscle disorders (*6*). We now show that the actomyosin cortex, regulated by Arpc5-containing Arp2/3 complexes, acts as a gatekeeper for membrane availability required for t-tubule growth. This is essential for ensuring proper myofiber development and muscle function and provides a novel pathophysiological mechanism for muscle disorders with impaired triad function.

Our observations reveal that the destabilization of the actomyosin cortex releases plasma membrane, enabling t-tubule extension. Membrane tubulation has also been observed following stretch release in adherent cells, suggesting that membrane release facilitates tubule formation (*39*). In addition, elegant experiments using giant unilamellar vesicles have demonstrated that membrane release alone, induced by a reduction in vesicle volume, can drive inward tubulation, mimicking t-tubule formation without the need for external forces (*40*). Importantly, we did not observe enrichment of actin or accumulation of branched actin near growing t-tubules. While we cannot exclude the involvement of external forces, our data does not support a requirement for actin-mediated force generation in t-tubule growth. Bin1 has also been shown to induce membrane tubulation independently of actin (*41*). Bin1, along with proteins such as dynamin-2, likely contributes to t-tubule growth by stabilizing membrane curvature, in synergy with the mechanism we identify (*41*, *42*).

Our findings support a model in which limiting t-tubule growth is essential for proper triad formation and synchronized muscle contraction. Excessive t-tubule growth may disrupt membrane contacts with the endoplasmic reticulum, impairing triad assembly and function. Postnatal loss of Arpc5 results in t-tubule abnormalities that might lead to the observed muscle fatigue, highlighting its role in maintaining t-tubule architecture. During postnatal development, t-tubules and triads undergo spatial consolidation into a transverse orientation and our results suggest that actomyosin cortex regulation has a role in this process (*7*, *36*). T-tubule overgrowth and triad abnormalities have been observed in aged human muscle (*43*). Moreover, several muscle disorders characterized by defective triads, including those involving mutations in Mitsugumin 25, Dysferlin, Myotubularin, and Dynamin-2, also exhibit excessive t-tubule clustering (*3*, *6*). Thus, the mechanism that regulates t-tubule growth that we identify may be a common therapeutic target for these disorders.

## Supporting information

Vide S9

## Acknowledgments

We thank all the members of the Gomes, Way and Moores laboratories for their continued support, scientific discussions and helpful comments on this work. We thank A.L. Sousa from the Electron Microscopy Facility at the Gulbenkian Institute for Molecular Medicine for her technical expertise, sample processing, and imaging of differentiated myofibers and postnatal myofibers by transmission electron microscopy. We also thank the Bioimaging Unit (funded by PPBI-POCI-01-0145-FEDER-022122) and the Rodent Facility of the Gulbenkian Institute for Molecular Medicine for their technical support. We thank N. Lukoyanova and S. Chen for electron microscope support and D. Houldershaw for computational support at Birkbeck. We finally thank the Molecular and Transgenic Tools Platform (MTTP) of Champalimaud Foundation for cloning and AAV production.

## Author contributions

A.R.P. and A.R.S. carried out experiments and analysed data. T.L., J.L.F. and C.A.M designed the cryo-CLEM experiments, T.L and J.L.F. collected the cryo-ET data, T.L. performed the 3D reconstruction and segmentation the experiments and, T.L and C.A.M analysed the data. S.d.F. performed experiments with Arpc5 KO mice and analysed data. G.L. performed the t-tubule cluster imaging. I.F. generated GFP-Bin1 lentivirus expressing plasmid. F.A.C. and N.C.S. carried out AFM experiments. N.K. and M.W. provided unpublished data, tools, and animal models. A.R.P, A.R.S, and E.R.G designed experiments and wrote the manuscript with assistance from other authors. All authors participated in the critical review and revision of the manuscript. E.R.G, M.W and C.A.M supervised the project and secured funding.

## Data and materials availability

All data are available in the main text or the supplementary materials.

**Supplementary Materials** Materials and Methods Supplementary Text Figs. S1 to S5 Movies S1 to S18

## Materials and Methods

### T-tubule labelling in mature myofibers

All procedures using animals were approved by the Institutional Ethics Committee and followed the guidelines of the Portuguese National Research Council Guide (DGAV). Ex vivo isolated single myofibers from Extensor Digitorum Longus (EDL) muscles were generated using the protocol previously described (*34*). EDL muscle was explanted from 8- to 12-week-old male or female C57BL/6J mice and then digested in Dulbecco’s modified Eagle’s medium (DMEM, Invitrogen) containing 0.2% (w/v) type I collagenase (Sigma) for 2 h at 37°C. Mechanical dissociation of fibers was performed using a thin Pasteur pipette in a transilluminating-fluorescent stereomicroscope. Single fibers were collected and plated in ibidi-35mm-dishes pre-coated with 1% (v/v) matrigel in Iscove’s Modified Dulbecco’s Media (IMDM, Invitrogen) with Glutamax. After plating, cells were covered with an upper coating of ice-cold 50% (v/v) matrigel coating in IMDM with Glutamax. Plated myofibers were maintained in 10% (v/v) horse serum (Fisher Scientific) plus IMDM with Glutamax for the duration of the experiment. To visualize t-tubule dynamics, plated myofibers were labelled for 30 min with CellMask Orange (1 µM, Invitrogen).

### In vitro myofibers

All animal procedures were approved by the institutional ethics committee and followed the guidelines of the National Research Council Guide for the care and use of laboratory animals. In vitro myofibers were differentiated as previously described (*44*) from primary myoblasts isolated from 5 to 7-day-old mixed-gender C57BL/6 mice. Tibialis anterior (TA), EDL, gastrocnemius and quadriceps muscles were dissected and kept in ice-cold PBS. Muscles were digested mechanically using surgical scissors and incubated for 90 min at 37⁰C with agitation in collagenase type V (Sigma) and dispase II (Roche) in DPBS. Digestion was stopped by mixing 10% (v/v) of fetal bovine serum (FBS) plus 1% (v/v) of Penicillin-Streptomycin IMDM with Glutamax. The cells suspension was centrifuged for 5 min at 75 x g to remove debris followed by a second centrifugation for 5 min at 350 x g. After centrifugation, cells were resuspended in the same medium and pre-plated for 4 hours to allow for fibroblasts adherence. The cells remaining in suspension were collected, centrifuged for 5 min at 350 x g and plated at a density of 220 000 cells/ml in 20% (v/v) FBS plus IMDM with Glutamax in ibidi 35 mm dishes pre-coated with 1% (v/v) matrigel in IMDM with Glutamax. After three days of proliferation, medium was replaced by 10% (v/v) horse serum IMDM with Glutamax supplemented with 100 ng/ml recombinant agrin to induce differentiation. An ice-cold upper coating of 50% (v/v) matrigel in 10% (v/v) horse serum IMDM with Glutamax was added the following day and fresh medium was added. Experiments were performed either at 4-5 or 7-8 days after differentiation.

### Mice

Skeletal muscle-specific Arp2/3 knockout mice strains mice were developed by breeding Arpc5 floxed mice (*37*) with mice expressing Cre recombinase (MerCreMer or MCM) double fusion protein under the human α-skeletal actin (HSA) promoter (*38*). The HSA promoter drives the expression of a tamoxifen (TMX)-inducible specifically in skeletal muscle. Upon tamoxifen administration, the Cre recombinase will be translocated in the nucleus where it will excise the Arpc5 floxed gene only in myofibers (Fig. 4A). For each experiment, cre-negative littermates were used as controls (Arpc5^WT^). Both males and females were used without distinction. 4-Hydroxytamoxifen (Sigma, H6278) was injected intraperitoneally (40 µl of 1,3 mg/mL solution) at postnatal (P) days 1, 2 and 3. At P7-P8, motor function was evaluated and/or animals were euthanized by decapitation. Hindlimb muscle was dissected for histological analysis and electron microscopy processing. All the procedures were approved by the Institutional ethics committee and followed the guidelines of the Portuguese National Research Council Guide (DGAV Project Licenses n° 022871/2016 and 004560/2022) for the care and use of laboratory animals.

### Motor function evaluation

To evaluate motor function postnatally we used a test appropriate to neonates, namely the hindlimb suspension test (*45*). Hindlimb suspension test; pups were placed gently facing down into a 50 mL conical tube, with their hind legs hung over the rim. This was repeated 3 times, with approximately 30 s of resting time among each trial. Finally, fatigue was assessed by calculating the ratio between the time to fall in the final trial and the initial trial. This fatigue index (final time / initial time) reflects the decline in endurance capacity over repeated tests. A lower value indicates a greater degree of fatigue, as a reduced ability to sustain suspension over time.

### siRNAs, plasmids, and transfections

To image F-actin, GFP-Bin1 myoblasts were transfected with 1 µg plasmid/ibidi-35mm-dish of LifeAct-mCherry (*46*) or mScarlet-I-UtrCH (gift from Dorus Gadella, Addgene plasmid #98823; http://n2t.net/addgene:98823; RRID:Addgene_98823 (*47*)) one day after plating, using Lipofectamine 2000 (Thermo), according to manufacturer’s protocol. For Arpc5 depletion, GFP-Bin1 myoblasts at 30% confluency were transfected with Thermo Fisher Silencer Select siRNAs for Arpc5. Two different siRNAs for Arpc5 were used: #1 (ID#s85623) and #2 (ID#s85621) at 20nM/ibidi-35mm-dish using Lipofectamine RNAiMAX (Thermo). Silencer® Select Scramble (ID #4390843) was used as a negative control. Cells were imaged by vtiSIM microscope on the next day (see below).

Triadin-GFP (Trisk95-GFP, (*48*), WT-Ezrin (Ezrin-GFP, gift from Guillaume Charras lab) or CA-Ezrin (EzrinT567D-GFP, gift from Guillaume Charras lab) plasmids were transfected to in vitro myofibers three days after plating (differentiation day 0) with 1 µg of DNA per ibidi-35mm-dish of the appropriate plasmid using Lipofectamine 2000 (Thermo), according to manufacturer’s protocol. To deplete Arpc5 in myofibers, Thermo Fisher Silencer Select siRNAs for Arpc5 #1 (ID#s85623) was transfected 3 days after plating at 20nM/ibidi-35mm-dish using Lipofectamine RNAiMAX (Thermo). Silencer Select Scramble (ID #4390843) was used as a negative control.

### Generating the GFP-Bin1 myoblast stable cell line

GFP-Bin1 stable cell line of C2C12 myoblasts (GFP-Bin1 myoblasts) was generated using a pLVX-puromycin vector containing Bin1 isoform 8. The lentiviral vector was constructed with VectorBuilder. The vector ID is VB220728-1365nbr, which can be used to retrieve detailed information about the vector on vectorbuilder.com. Lentivirus were produced in HEK 293T cells. To generate stable cell lines expressing exogenous GFP-Bin1, C2C12 myoblasts were plated in a 6-well plate. The cells with a 10% confluency were carefully covered with 0.5 mL of filtered medium containing the lentiviruses and 0.5 mL of complete medium was added to myoblasts (1 mL total volume). The medium containing lentiviruses was left over day, then 1 mL of complete fresh medium was added. Cells were allowed to grow until 80% of confluency. Subsequently, the sorted myoblasts were allowed to expand, and the process was repeated for three rounds of sorting to achieve a homogeneous cell population.

GFP-Bin1 myoblasts were cultured in DMEM with glutamine (Gibco) supplemented with 10% FBS (Gibco) and penicillin-streptomycin (100 U/mL:100 μg/mL; Gibco), at 37°C under 5% CO_2_.

### AAV1-CAG-CAAX-GFP production

The AAV1-CAG-GFP-CAAX was generated following the protocols from Addgene with minor modifications (*49*, *50*). Briefly, production began by triple transfecting the pAAV-CAG-CAAX plasmid (see Supplementary Text for plasmid map) with AAV packaging plasmid expressing Rep2 and Cap1 (#112862, Addgene) and pHelper into 15-cm plates of confluent HEK-293T cells using polyethylenimine (Polysciences Inc., catalog # 23966-100). The cells were harvested three days post-transfection. The purification step was performed by iodixanol gradient ultracentrifugation and the viral fraction concentrated to approximately 150 μL using an Amicon Ultra-15 centrifugal filter unit (Merck, catalog #UFC910024). For measuring virus titer, we performed qPCR (Biorad CFX96 Real-Time PCR system), using primers against Woodchuck Hepatitis Virus Posttranscriptional Regulatory Element (wpre).

### Fluorescence microscopy

Airyscan images were acquired using a Zeiss LSM980 Airyscan2 microscope, equipped with a 63x Plan-Apochromat DIC oil objective (NA 1.4), unless stated otherwise. Imaging of fixed cells was performed at RT, and imaging of live cells was performed at 37°C with 5% of CO_2_ using the ZEN 3.8 blue edition. To measure t-tubule growth, images were recorded every 3 sec. Airyscan reconstruction was performed using automatic filter settings. To visualize the t-tubule network of in vitro differentiated myofibers at higher resolution, Airyscan images were processed using joint-deconvolution (jDCV) in ZEN blue, which is an accelerated joint Richardson-Lucy algorithm (Zeiss).

For instant structured illumination microscopy (iSIM) imaging, the vtiSIM module (Visitech) equipped on a Nikon Tie microscope with a 100x AC Oil CFI SR HP Apo TIRF objective, and a Photometrics Prime BSI Express Scientific sCMOS camera was used. Timelapse imaging was performed at 37°C with 5% CO_2_.

To visualize t-tubule dynamics in developing myofibers, the membrane lipid dye CellMask Orange (1 µM, Invitrogen) was added to 4 days differentiated myofibers 5-10 min before imaging. T-tubule and/or Triadin-GFP dynamics were recorded by timelapse imaging using Airyscan microscopy (see above).

To visualize actin dynamics near growing t-tubules, GFP-Bin1 myoblasts transfected with LifeAct-mCherry or mScarlet-I-UtrCH were imaged by vtiSIM timelapse every sec for 3 min.

To visualize t-tubule network and t-tubule clusters, developed in vitro myofibers (differentiation day 7) were incubated with 1 µM CellMask Deep red (Invitrogen). Spinning disk confocal imaging was performed using a Zeiss Axio Observer microscope equipped with a Yokogawa CSU-X1 scanning unit, an Evolve 512 EMCCD camera and a 63x Plan-Apochromat DIC oil objective (N.A. 1.4), using the ZEN 3.2 blue edition software. Cells were kept at 37°C and 5% CO_2_. Myofibers with more than three t-tubule clusters were visually quantified using Fiji software (*51*) and the percentage of cells with t-tubule clusters was plotted using GraphPad.

### Drug treatments

Cells were treated with 100 µM of CK666 (Sigma) for Arp2/3 inhibition. A final concentration of 125 µM of para-aminoblebbistatin (Motorpharma) was used to block myosin-II. To inhibit Ezrin phosphorylation, NSC668394 (MedChemExpress) was used at 50 µM. The number of growing tubules or t-tubules was assessed immediately after drug treatments for GFP-Bin1 myoblasts and after 5 min for myofibers. T-tubule and tubule dynamics were recorded by timelapse imaging using Airyscan microscopy. GFP-Bin1 myoblasts treated with para-aminoblebbistatin or glycerol were recorded by timelapse imaging using vtiSIM microscopy.

### Glycerol treatment

Myoblasts and developing myofibers were exposed to 225 mM and 45 mM of glycerol, respectively, to induce hyperosmotic shock. The number of growing t-tubules was performed 3 min after glycerol treatment for GFP-Bin1 myoblasts and after 5 min for myofibers. Myofiber t-tubule dynamics were recorded by timelapse imaging using Airyscan microscopy. GFP-Bin1 myoblasts t-tubule dynamics were recorded by timelapse imaging using vtiSIM microscopy.

### Immunostainings

Developing myofibers (differentiation day 4) or differentiated myofibers (differentiation day 7) were fixed in 4% paraformaldehyde in PBS for 10 min at room temperature and permeabilized with 0.5% Triton X-100 for 5 min. Cells were blocked in a solution containing 10% goat serum, 5% BSA and VisUBlock Mouse on Mouse Blocking Reagent (R&D Systems, VB001). Primary and secondary antibodies were diluted in 0.1% saponin. Cells were incubated with primary antibodies overnight at 4°C and washed three times for 10 min with PBS. Cells were incubated with secondary antibodies for 1 h at room temperature and washed three times for 10 min with PBS. Cells were mounted with Fluoromount-G (Invitrogen).

### Junctophilin1 immunostaining

For Junctophilin1 immunostaining in differentiated myofibers (day 7), we used an anti-JPH1 antibody (1:200 dilution, Invitrogen, 40-5100) and an anti-rabbit Alexa Fluor 488 secondary antibody (1:400, Life Technologies). Phalloidin conjugated with an Alexa Fluor 647 (1:200, Life Technologies) was used to stain filamentous actin. Myofibers were imaged using the vtiSIM microscope (VisiTech).

### p-ERM and NM-myo2B localization at the plasma membrane

To label the plasma membrane, in vitro primary myofibers were infected on day 2 of differentiation with the AAV1-CAG-GFP-CAAX (1.535 x 10^^7^ genome copies to a 35mm dish containing myofibers at approximately 70-80% confluency). The virus was washed on the next day, with differentiation media containing 1 µg/ml of agrin. At day 4 of differentiation, cells were fixed and stained as described above.

To label pERM we used an anti-Phospho-Ezrin (Thr567)/Radixin (Thr564)/Moesin (Thr558) (1:200 dilution, Cell Signaling Technology #3141) and anti-rabbit Alexa Fluor 555 secondary antibody (1:400, Life Technologies). To label NM-myo2B, cells were incubated with an anti-Non-muscle Myosin Heavy Chain II-B (NM-myo2B) antibody (1:200 dilution, Biolegend) and anti-rabbit Alexa Fluor 555 secondary antibody (1:400, Life Technologies). Cells were also co-stained with an anti-GFP antibody (1:200 dilution, Roche #11814460001) and anti-mouse Alexa Fluor 488 secondary antibody (1:400, Life Technologies). Cells were imaged by Airyscan using the LSM980 microscope with a 40x LD C-Apochromat, N.A. 1.1, Water immersion objective.

To quantify the number of foci of NM-myo2B, using Fiji, we drew rectangle ROIs with a constant width of 0.8 µm along the CAAX-GFP channel. The NM-Myo2B fluorescence intensity signal over that ROI was then plotted and the number of peaks (corresponding to NM-Myo2B foci) was measured with the Find Peaks BAR tool (*52*) using a mean amplitude of 100. The number of peaks in each ROI was divided by the respective area of the ROI, and these values were plotted as the number of NM-Myo2B foci per ROI area (µm^2^).

To quantify pERM at the plasma membrane, we used the CAAX-GFP signal to identify the plasma membrane and drew a line of 10 µm length by 10 pixels width on top of the signal, and extracted the single-pixel intensity values of pERM. To calculate the pERM intensity signal, we averaged the single-pixel intensity over the 10 µm line. To measure the pERM coefficient of variation, we calculated the standard deviation of the single-pixel intensity over the 10 µm line divided by the mean of fluorescence.

### Atomic force microscopy

Atomic force microscopy (AFM) imaging of developing myofibers was performed using a NanoWizard IV system (Bruker-JPK Instruments, Berlin, Germany) mounted atop an Axiovert 200 inverted optical microscope (Carl Zeiss, Jena, Germany). To avoid cell contraction during acquisition, developing myofibers (differentiated for 4 days) were imaged in Tyrode Buffer (140 mM NaCl, 5 mM KCl, 2 mM MgCl_2_ and 10 mM HEPES, pH 7.2). For AFM imaging, oxidized and sharpened silicon probes with a nominal tip radius of approximately 6 nm, a resonance frequency of 60 kHz, and a spring constant of 3 N/m were employed. Scanning was performed at an optimized speed of 0.3 Hz using a minimal applied force to avoid damaging the sample. Images were acquired at a resolution of 512 × 512 pixels.

Surface roughness measurements were conducted on live cells following four days of differentiation, using the same equipment. Experiments were performed at 25°C in Tyrode’s solution, employing the Quantitative Imaging (QI) mode to capture height data. Non-functionalized qp-BioAC CB2 cantilevers (Nanosensors, Neuchâtel, Switzerland) with partial gold coating and quartz-like tips, featuring a nominal spring constant of 100 pN/nm, were used. Differential interference contrast (DIC) microscopy facilitated accurate positioning of the cantilever tip over both Arpc5 KD and scramble control cells. QI-mode height maps of 5 µm × 5 µm were acquired with a vertical (z) range of 2 µm. Images were recorded at a resolution of 256 × 256 pixels with a pixel dwell time of 20 ms. During each approach, the cantilever-to-cell distance was dynamically adjusted to maintain a maximal applied force of 1 nN prior to tip retraction. For each experimental condition, a minimum of 12 height images were obtained, with all samples analyzed in triplicate. Image processing was performed using JPK Image Processing software (v. 8.0.144) to calculate average roughness (Ra) and root mean square roughness (Rq). For each height map, roughness was determined in five distinct 0.5 µm × 0.5 µm square regions located between Z-bands. In total, surface topographies of 12 control cells and 15 Arpc5 KD cells were analyzed.

### Image analysis

All images were processed in Fiji (*51*). For cell drift correction we used the Fiji plugin Fast4Dreg (*53*).

For t-tubule growth speed measurements, manual tracking of the tip of the t-tubule during the timelapse was performed using Fiji manual tracking.

The number of growing t-tubules was manually recorded using the Fiji cell counter tool. This value was then normalized for the cell area. All quantifications were performed within 2 min timelapses (42 frames). For the cells expressing WT-Ezrin or CA-Ezrin, the number of growing t-tubules was measured in cells containing GFP positive signal only.

To measure Triadin-GFP mean speed in developing myofibers, we used Airyscan images of Triadin-GFP timelapses acquired every 3 sec for 2 min. Since Triadin-GFP localizes as distinct spots, we used the Fiji plugin Trackmate (*54*) to track Triadin-GFP spots over time in myofibers. To detect Triadin-GFP spots we used the LoG detector with an estimated object diameter of 0.5 μm and a quality threshold of 5, 10 or 20. We then linked the spots to generate tracks, by using the LAP tracker with a max distance of 0.5 μm and no gap closing. To have enough datapoints to calculate speed trajectories, we only considered tracks that lasted more than 10 frames (30 sec). The resulting tracked Triadin-GFP spots were then used to calculate mean speed.

To quantify actin dynamics in the vicinity of growing t-tubules, we manually selected a ROI around GFP-Bin1 growing tubules (as the ones represented in fig. S2A and S2C), including a few timepoints before the GFP-Bin1 signal was observed using Fiji software. We then extracted at each timepoint the average of fluorescence intensity per µm^2^ for GFP-Bin1, LifeAct-mcherry or mScarlet-I-UtrCH using the Fiji software. Values were normalized for the initial corresponding average intensity, and these normalized intensity values per µm^2^ over time were plotted using GraphPad.

To automatically count the number of Junctophilin 1 (JPH1) puncta, a maximum projection of the Z-stack images that were in focus was first performed using Fiji. Then, a rectangular ROI with an area of 175.552 μm^2^ was drawn along the myofiber length. To detect the puncta automatically, we used the LoG detector tool from the Fiji TrackMate plugin (*54*). LoG filters for puncta detection were a blob diameter of 0.5 μm and an intensity threshold of 5. To analyse the number of JPH1 puncta per µm^2^, the number of detected puncta was normalized for ROI area and plotted.

### Western blot

Cell extracts were collected in 100 mM Tris-HCl pH 8 with 1% (w/v) SDS. Tissue was extracted from snap-frozen muscle samples and homogenized using an Omni Soft Tissue Tip Homogenizer (Biolabproducts GmbH). Protein lysates were prepared using RIPA buffer (NaCl 150 mM, EDTA 5 mM, 1% (v/v) Triton X-100, 0.5% (v/v) NP-40, Tris 50 nM pH 7.5) with protease and phosphatase inhibitors (Roche). All protein quantification was performed using the BCA Protein Assay Kit (Pierce) following the manufacturer’s protocol. Samples were denatured in Laemmli buffer and heated at 95°C for 5 min. 10-30 ug of total protein was loaded onto a 10% (v/v) precast polyacrylamide gel (Bio-Rad) and transferred to a nitrocellulose membrane for 75 min at 100 V. The membranes were blocked with Tris-buffered saline with 0.1% (v/v) Tween 20 (TBS-T) containing 5% skim milk, followed by incubation with primary antibodies, overnight at 4°C. Membranes were washed and incubated with HRP-conjugated secondary antibodies for 1 hour at room temperature and washed three times before image acquisition. Images were acquired using Amersham ImageQuant800 (Cytiva). Antibodies used were: anti-Arpc5 (1:500 Santa Cruz Biotechnology, sc-166760); anti-Vinculin (1:1000, Sigma, V9131); anti-rabbit HRP (1:5000, Thermo Scientific) and anti-mouse HRP (1:5000, Thermo Scientific).

### Cryo-ET sample preparation

GFP-Bin1 myoblasts were grown to approximately 60% confluency and then diluted to a confluency of 10%. EM grids (Quantifoil R3.5/1 Au 200 mesh) were glow-discharged using a plasma cleaner (PLASMAFLO PDC-FMG-2, Harrick Plasma) and then placed in a 35mm MatTek dish in a flow hood. 300 μL of 5 μg/mL Fibronectin (Sigma-Aldrich, 341635) was added to the dish, covering the EM grids. The dish with the grids inside was then sterilized under UV light for 1.5 h. After simultaneous coating and sterilization, fibronectin was removed from the MatTek dish. The grids were then washed twice with PBS and stored in approximately 300 μl of medium (DMEM high glucose+pyruvate+L-glutamine; 10% (v/v) FBS + 1% (v/v) PenStrep).

Once the cells were ready, the medium in the grid-containing dish was removed, and diluted cells were added to the dish. The cells were seeded on the EM grids at 37°C 5% CO_2_ overnight (∼15 h). The following day, grids were inspected under an inverted biological microscope (Medline Scientific, 3650.0000) to check the cell density and ensure grid integrity. Grids with minimally broken areas and well-distributed cells were plunge frozen using Leica GP2 with back blotting for 15 s and an additional movement of 0.5 mm at 37°C and 90% humidity.

### Correlative light and electron microscopy

Frozen grids were first loaded onto a Titan Krios microscope (Thermo Fisher) operated at an accelerating voltage of 300 kV with a K3 detector and a BioQuantum energy filter (Gatan). Grid atlas images were collected to check the ice quality and the cell density using EPU software (Thermo Fisher). After locating the spread cells, grids were transferred onto a cryo-light microscope (Leica THUNDER Imager EM Cryo CLEM). Z stack fluorescence and bright field images were taken of the located spread cells using LAS X software at 50x magnification. With the fluorescence images processed using THUNDER technology, individual tubular fluorescence signals were observed. The fluorescence images were saved for fluorescence microscopy (FM) and electron microscopy (EM) image correlation. The grids were then transferred back to a Titan Krios microscope (Thermo Fisher) operated at an accelerating voltage of 300 kV, with Selectris X image filter (Thermo Fisher) and Falcon 4i camera (Thermo Fisher). Medium magnification atlases were taken and combined on each fluorescence-labelled cell for manual correlation and locating the regions of interest. Empty holes and dark features were used as markers to facilitate the manual correlation (fig. S3B and S3C). After correlation, ROI with fluorescence signals were targeted for cryo-ET tilt series data collection. 114 tilt series were collected using TOMO5 software with a dose symmetric tilt scheme +/- 60° with the step size of 3° at the magnification of ×42,000, pixel size of 3.0 Å, dose rate of 12.6 e^-^/pixel/s, exposure time of 2.69s and total dose per micrograph of 3.66 e^-^/ Å^2^.

### Tomogram reconstruction

Tomograms were reconstructed using RELION 4.1 tomography pipeline (*55*). Raw data were imported into RELION followed by motion correction with RELION’s own implementation of MotionCor2 (*56*) and CTF correction with ctffind 4.1 (*57*). Tilt images with significant contamination and grid bars were manually excluded from further data processing in a Napari-based viewer (Exclude Tilt Images job). Then each tilt series was aligned using IMOD _[CM4]_ patch tracking with Patch size 100 nm and Patch overlap 50%. The aligned tilt series were used to generate tomograms with a binned pixel size of 10 Å. Tomograms were denoised using CryoCARE to improve the signal-to-noise level (*58*). CryoCARE training was performed with 1200 sub-volumes per tomogram and a sub-volume dimension of 72 pixel. Upon inspecting the tomograms, we found that 58 contained vesicular as opposed to tubular membrane structures (as shown in fig. S3G) and were excluded. 23 of the tomograms were discarded due to the presence by non-vitreous ice and could not be used for segmentation. Additionally, 27 tomograms did not correlate well with fluorescence signals because of the ambiguity of fluorescence and the lack of correlation markers, which posed challenges for manual correlation. 6 tomograms displayed tubular structures at locations where fluorescence was observed. From these, the 3 highest quality tomograms were selected for segmentation and further analysis.

### Tomogram segmentation

Tomograms were processed using IsoNet to further improve the signal-to-noise (snr) level (*59*). The following parameters were used: CTD deconvolve snrfalloff of 0.5, mask making density_percentage of 100, std_percentage of 100, and z_crop of 0.17. Then, IsoNet-processed tomograms were cropped to remove regions lacking t-tubule structures to simplify segmentation. Segmentation was performed using the neural network-based software Dragonfly (Dragonfly 2024.1 [Computer software]. Comet Technologies Canada Inc., Montreal, Canada; software available at https://dragonfly.comet.tech/). For each tomogram, actin and membrane structures were manually annotated on 5-12 sub-areas across different slices of the tomogram. The segmented data were then used to train multiple deep-learning models. The model segmenting the highest number of actin filaments on test slices was selected to segment the entire tomogram. The UNet++ model (2.5D, 3 slices, depth level of 5, and initial filter count of 32) was used for tomograms 1 and 2. Additionally, a pretrained UNet model provided by the Dragonfly team (depth level of 5 and initial filter count of 64) was utilized for tomogram 3. The segmentations were manually cleaned and corrected in Dragonfly. The segmented actin and membrane structures were rendered into a 3D contour mesh and smoothed over five to ten iterations for 3D visualization in fig. S3I and fig. S4. Movies were created by exporting the segmented actin and membrane structures as binary files, converting these into two MRC files, and then importing them along with a contrast-inverted 3D tomogram into ChimeraX for visualization (*60*).

### Tomogram analysis

For tomogram analysis, sub-tomograms proximal to the t-tubules and distant from the t-tubules were selected. The sub-volumes near t-tubules were constrained to regions no more than 200 nm from the t-tubule structures. Actin branches – inferred from the relative position of two connected filaments - within these boxes were identified, labelled, and counted. For control purposes, sub-volumes located more than 200 nm away from t-tubules were also identified, and suspected actin branches within these control sub-volumes were similarly labelled and counted. In total, 7 sub-volumes were selected to determine the number of actin branches near the t-tubules, and 3 sub-volumes were selected as controls.

### Single-myofiber contraction and calcium transient assays

In vitro myofibers were plated on 35mm fluorodishes (World Precision Instruments). After 7-8 days of differentiation, developed myofibers were imaged using a Nikon Eclipse Ti microscope. Myofibers were kept at 37°C and CO_2_ 5% using an Okolab stage incubator throughout all experimental procedures. Electrical field stimulation using the MyoPacer system (Ionoptix) was used to induce excitation-contraction of myofibers using 10 V at 0.5 Hz frequency pulses with a duration of 20 ms.

To measure cytosolic calcium transients during electrical pulses, myofibers at differentiation day 3-4 were infected using 1 μl/mL of the AAV9-pAAV.CAG.GCaMP6s.WPRE.SV40 (Addgene viral prep # 100844-AAV9; the plasmid was a gift from Douglas Kim & GENIE Project (*61*)). After overnight incubation, the media was replaced with differentiation media (IMDM with Glutamax supplemented with 100 ng/mL recombinant agrin). After 7-8 days of differentiation, GCaMP fluorescence during electrical field stimulation was recorded using a 60x Plan Apo DM Ph3 (NA 1.4) objective, and an Andor Sona CSC-00324 camera at a 20 ms frame rate with the NIS-Elements AR 5.42.05 software. Fiji software was then used to generate subcellular GCaMP fluorescence kymographs by drawing a 475.7 μm^2^ square ROI along the myofiber that was then was maximum projected, generating GCaMP fluorescence kymographs. The number of cells that presented GCaMP transients synchronous with the electrical pulse was then quantified from these kymographs, and plotted.

For single-myofiber contraction tracing, electrical field stimulation was performed as mentioned above. Individual cell contractions were recorded with a MyoCam-S3 camera (Ionoptix) and a 40x CFI Plan Apochromat Lambda D objective (NA 0.95), using the IonWizard 7.7.3.180 software. Contraction kinetics were based on CytoMotion (Ionoptix) Pixel Correlation changes relative to the reference frame taken at rest (i.e. before electrical pulses). To generate single-myofiber contraction tracings, pixel correlation (a.u.) curves were plotted using the Cytosolver Cloud (Ionoptix). Myofibers that during the electrical stimuli did not contract, contract, present synchronous or asynchronous contraction during the electrical stimulus were then quantified and plotted.

### Transmission electron microscopy

Developed myofibers (7 days of differentiation) were fixed for 1 h at 4 °C in 0.1 M Sorensen’s Phosphate Buffer pH 7.4 containing 2.5% (v/v) glutaraldehyde (Science Services EMS), 2% (v/v) formaldehyde (Science Services EMS) and 50 mM calcium chloride (Sigma). Samples were postfixed for 1 h at 4°C in the dark with 0.1 M Sorensen’s Phosphate Buffer pH 7.4 containing 1% (w/v) osmium tetroxide (Science Services EMS) and 0.8% (w/v) potassium ferrocyanide (Sigma-Aldrich). After five washes with distilled water, samples were stained for 1h at room temperature with 0.5% (w/v) uranyl acetate (Analar). Dehydration was performed using an ethanol gradient (30% 10 min room temperature, 50% 10 min room temperature, 75% overnight 4°C, 90% 10 min room temperature, and 100% three times 10 min each room temperature). Samples were embedded in Embed-812 epoxy resin (Science Services EMS) and hardened at 60 °C for at least 24 h. Ultrathin sections (70 nm) were obtained and collected in homemade formvar-carbon coated copper-palladium slot grids and counter-stained with 1% (w/v) uranyl acetate in 70% methanol and Reynolds lead citrate, for 5 min each.

For muscle tissue from mice, longitudinal sections of postnatal myofibers from mouse neonates were generated from hindlimb muscles. For this, after sacrifice, the skin was removed and the whole mice leg was incubated with fixative (0.1 M Sorensen’s Phosphate Buffer pH 7.4 containing 2.5% (v/v) glutaraldehyde (Science Services EMS), 2% (v/v) formaldehyde (Science Services EMS) and 50 mM of calcium chloride (Sigma)) for 1 h, to preserve muscle structure. Rinse in 0.1M cacodylate buffer and dissecting by cutting small longitudinal pieces (<1mm3). These small pieces were incubated overnight at 4°C with fixative 0.1 M Sorensen’s Phosphate Buffer pH 7.4 containing 2.5% (v/v) glutaraldehyde (Science Services EMS), 2% (v/v) formaldehyde (Science Services EMS) and 50 mM calcium chloride (Sigma). Samples were postfixed for 2 h at 4°C in the dark with 0.1 M Sorensen’s Phosphate Buffer pH 7.4 containing 2% (w/v) osmium tetroxide (Science Services EMS) and 0.8% (w/v) potassium ferrocyanide (Sigma-Aldrich). After five washes with distilled water, samples were stained for 1h at room temperature with 0.5% (w/v) uranyl acetate (Analar). Dehydration was performed using an ethanol gradient (30% 10 min room temperature, 50% 10 min room temperature, 75% overnight 4°C, 90% 10 min room temperature, and 100% three times 10 min each room temperature). Samples were embedded in Embed-812 epoxy resin (Science Services EMS) and hardened at 60 °C for at least 24 hours. Ultrathin sections (70 nm) were obtained and collected in homemade formvar-carbon coated copper-palladium slot grids and counter-stained with 2% (w/v) uranyl acetate in 70% MetOH and Reynolds lead citrate, for 5 min each.

Regions of interest were mapped with SerialEM software and tiles of differentiated cells were acquired using a FEI Tecnai G2 Spirit BioTWIN (120 kV) transmission electron microscope with an Olympus-SIS Veleta CCD Camera.

Stitching was performed using the IMOD blendemont function (*62*). Stitched tiles were opened in Fiji and we counted the number of triads (one t-tubule connected to two ER) in each tile, and normalized it to the cell area. To count t-tubules or measure their area in myofibers, we identified the membrane objects that had a low electron-dense lumen, and that were contacting the ER (green or blue arrowheads in Fig. 3E and white arrowheads in Fig. 4). Area was measured by drawing a region of interest along the t-tubule membrane, using the drawing tool from Fiji software. We quantified membrane irregularity ratio as a measure of surface roughness. The irregularity ratio was calculated by dividing the length of the manually traced membrane contour by the length of a straight horizontal reference line drawn between the two ends of the membrane segment. Membrane outlines were manually traced using high-resolution TEM images in Fiji software. Values equal to one correspond to a flat membrane. Values higher than one correspond to the degree of membrane irregularity/roughness.

## Data representation and statistics

All graphs were generated and statistical analyses were conducted using GraphPad Prism v8. Normality testing was carried out using the Shapiro-Wilk test, with an alpha level=0.05. Details about the tests used are described in the Figure legends.

## Supplementary Text

### pAAV-CAG-GFP-CAAX_VectorMap

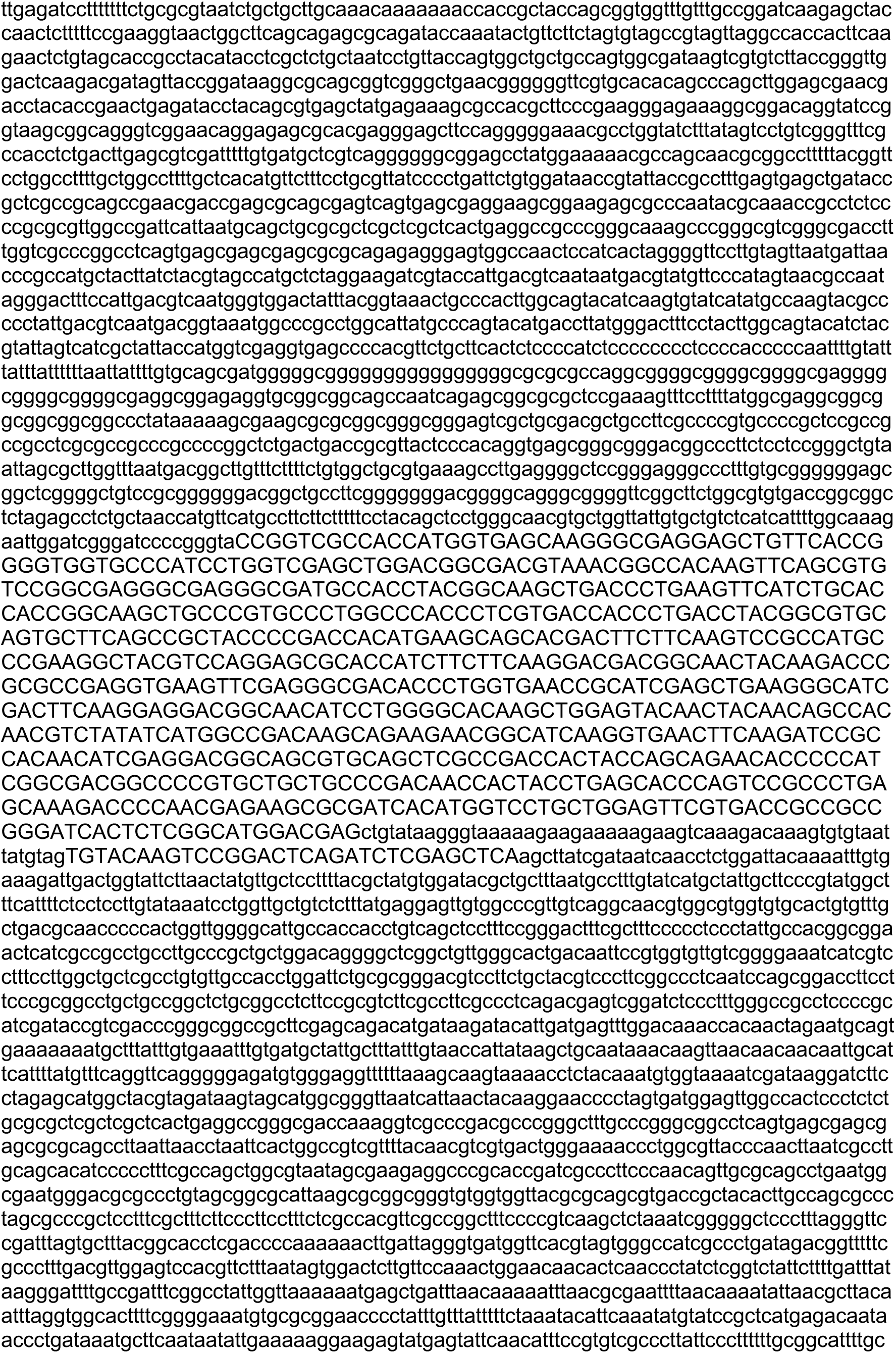
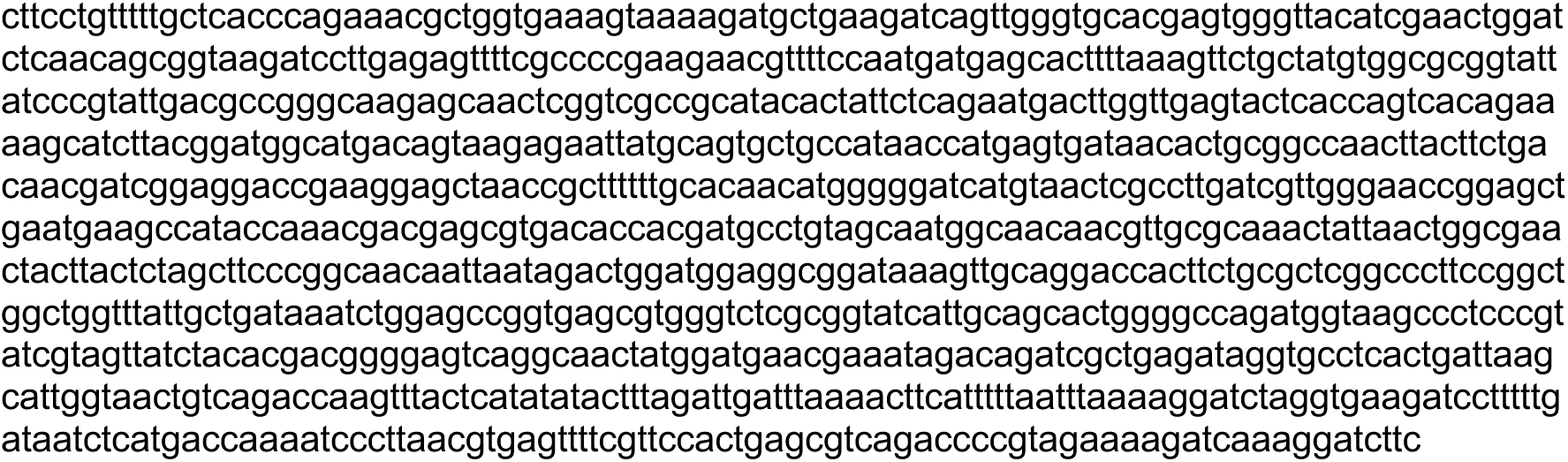

## Supplementary Figures

**Fig. S1.**
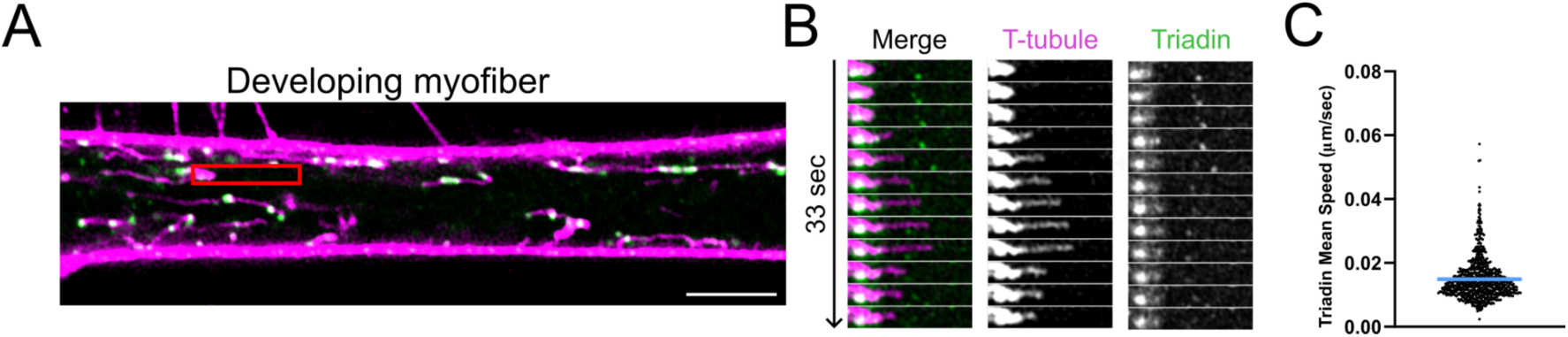
T-tubules are dynamic independently of contacts with the ER. Triadin-GFP, an ER transmembrane protein localises at t-tubule - ER contact sites and triads (*63*). (**A**) Timelapse snapshot of an in vitro developing myofiber (differentiation day 4) expressing Triadin-GFP (green), and CellMask DeepRed (magenta) to label t-tubules, was imaged by timelapse every 3 s for 2 min. Scale bar 5 µm. (**B**) Kymograph from red ROI showing that Triadin-GFP is static when t-tubules are growing. In 100% of growing t-tubules, we did not observe Triadin-GFP moving with the t-tubule (N=39 growing t-tubules). (**C**) The plot represents Triadin-GFP mean speed, calculated using trackmate, in five different cells from two independent repeats (N=783 tracks). Triadin-GFP exhibits a negligible average speed of 0.015 µm/s compared to the t-tubule growing average speed of 1.2 µm/s (Fig. 1B). Blue line on plot represents mean value.

**Fig. S2.**
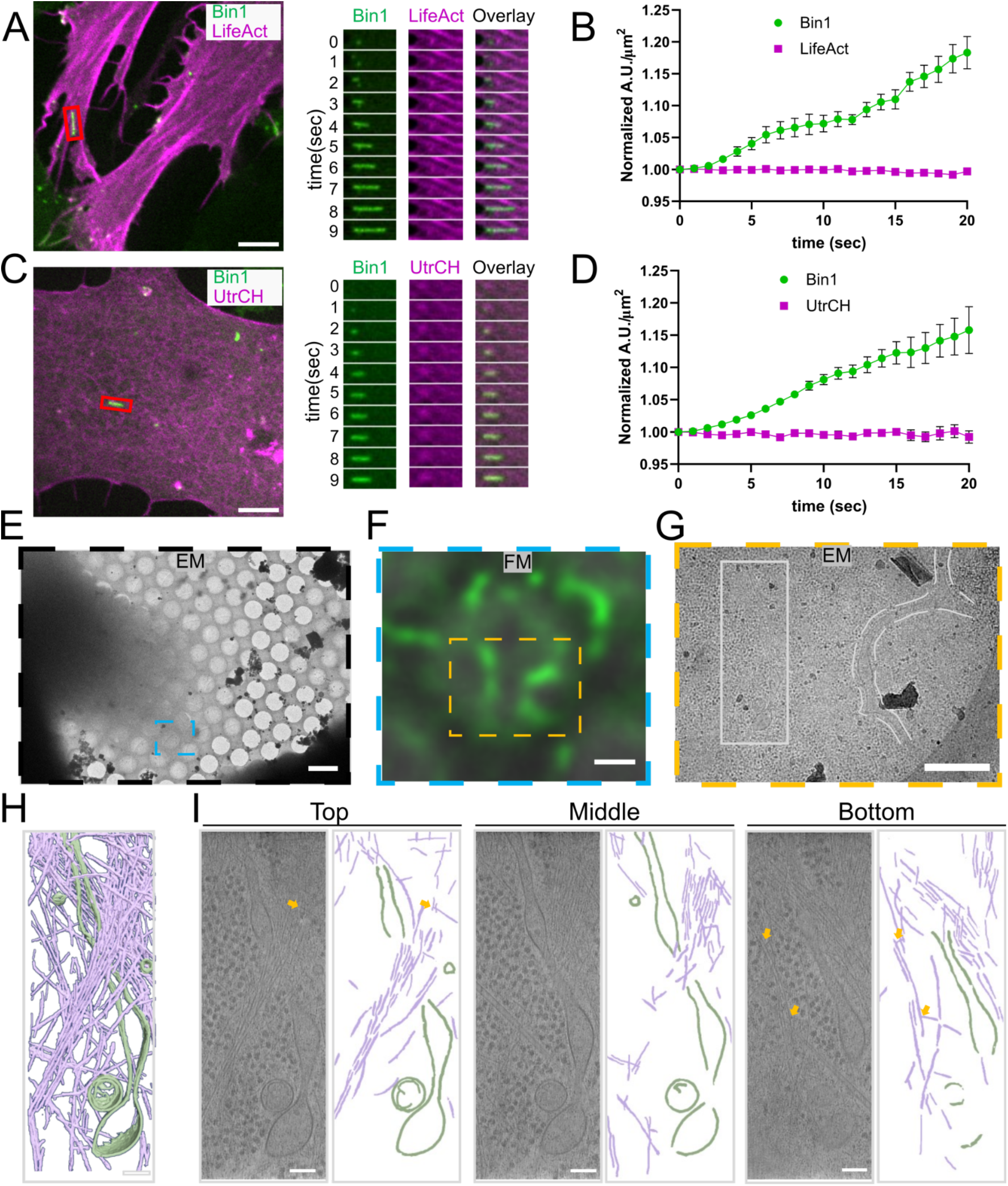
F-actin is not locally nucleated at growing tubules. **(A)** Left image corresponds to a timelapse snapshot of GFP-Bin1 myoblasts (green) and LifeAct-mCherry (magenta). Right image corresponds to kymographs from the red rectangle ROI showing a Bin1-tubule growing over time in the absence of F-actin changes. (**B**) Plots of LifeAct-mCherry and GFP-Bin1 average fluorescence intensity over time. Mean and SEM are represented. N=19 from two independent experiments. (**C**) Left image corresponds to a timelapse snapshot of GFP-Bin1 myoblasts (green) and mScarlet-I-UtrCH (magenta). Right image corresponds to kymographs from the red rectangle ROI showing a tubule growing over time in the absence of F-actin changes. (**D**) Plots of the mScarlet-I-UtrCH and GFP-Bin1 average fluorescence intensity over time. Mean and SEM are represented. N=16 from two independent experiments. (**E**) Representative low magnification cryo-EM image showing part of one cell on a TEM grid. The cyan rectangle indicates the ROI imaged using epifluorescence and brightfield microscopy (panel F). (**F**) Merged epifluorescence (green) and brightfield (grey, FM) image of the ROI. in E, showing multiple GFP-Bin1 tubules at the cell periphery. The orange rectangle indicates the ROI imaged using TEM (panel G). (**G**) Medium magnification TEM image of the ROI in F, showing the tubules observed in F. (**H**) 3D representation of cropped tomogram segmentation of the white rectangle ROI from G, with actin filaments (purple) and tubule membrane (green). (**I**) Slices of the tomogram in H at different z positions (from left to right: z=-25nm, 30nm and 135nm relative to the central slice) with actin filaments (purple), tubule membrane (green), and potential actin branches (yellow arrows). Scale bar: 5 μm in A, C and E, 1 μm in F, 500 nm in G and 100 nm in H and I.

**Fig. S3.**
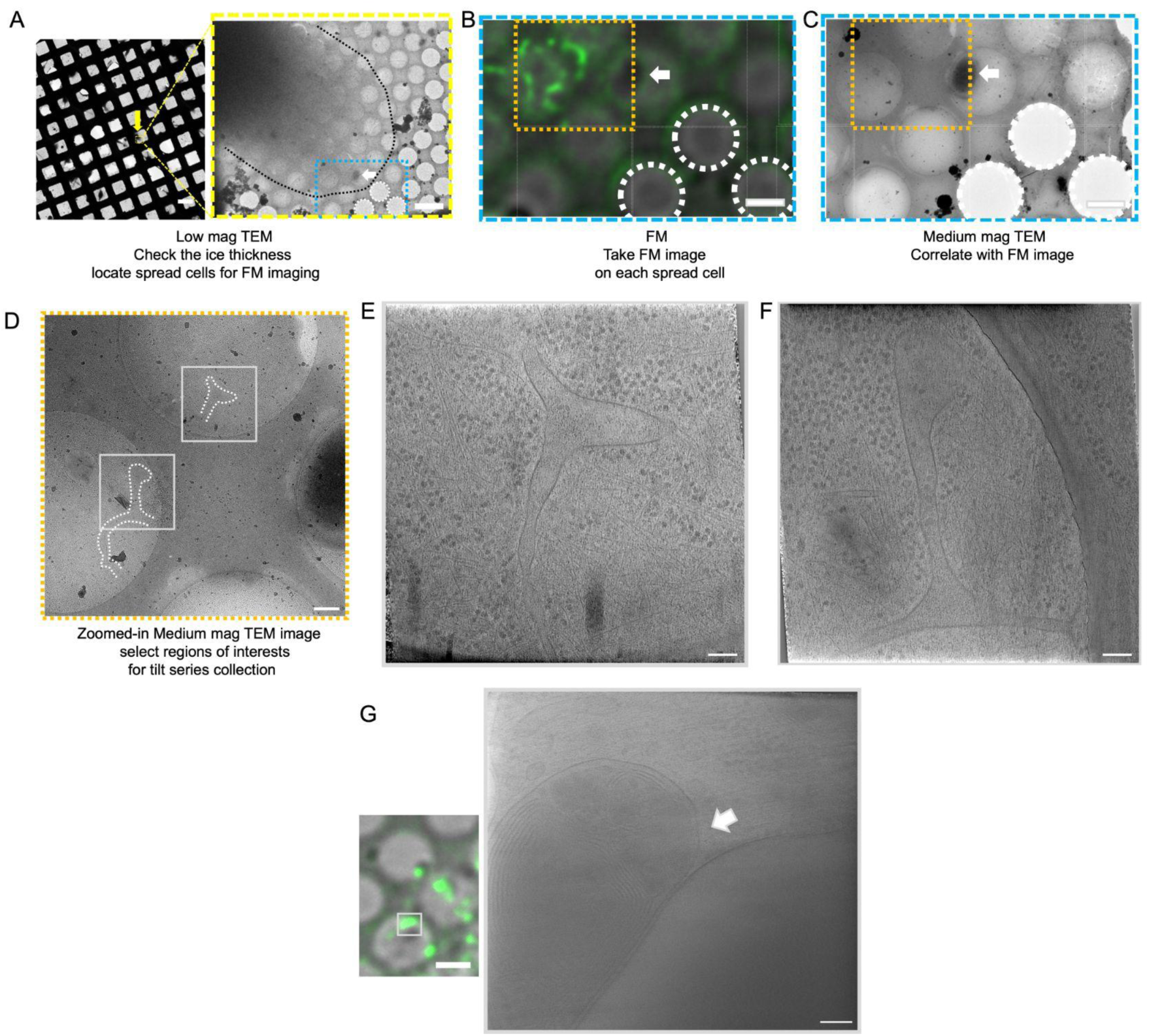
Correlative light and electron microscopy workflow. (**A**) Cells were imaged using TEM to check the ice thickness and locate spread cells for FM imaging. In the example image, one spread cell is indicated with a yellow arrow, with the approximate cell periphery outlined in black in the zoomed view. (**B**) Cells imaged using cryo-fluorescence microscopy (FM) with the bright-field image of the area in the blue box from A showing the GFP-Bin1-labeled membrane tubule structures at the cell periphery. (**C**) Cells were re-imaged using medium magnification TEM images in tiles and combined for FM-TEM correlation. The empty holes and black features were used for manual correlation, with empty holes indicated by white circles and the dark feature marked by a white arrow. (**D**) A zoomed-in view of the medium magnification image collected from the orange region in C. Tubular membrane structures are highlighted with white curvy lines. The shapes of these tubular membrane structures resemble those seen in the FM image in B. Regions of interest (ROI) were selected for tilt series data collection, indicated by white boxes. (**E**) and (**F**) The central slice along the z-axis of the 3D tomogram at positions 1 and 2, respectively. (**G**) The bright-field and FM image showing fluorescently-labelled vesicular structure. A tomographic slice of the reconstruction inside the white box showing nested sets of membranes is shown on the right. Scale bar: 100 μm and 5 μm in A, 2 μm in B and C,500 nm in D, 100 nm in E and F. 2 μm and 100 nm in G.

**Fig. S4.**
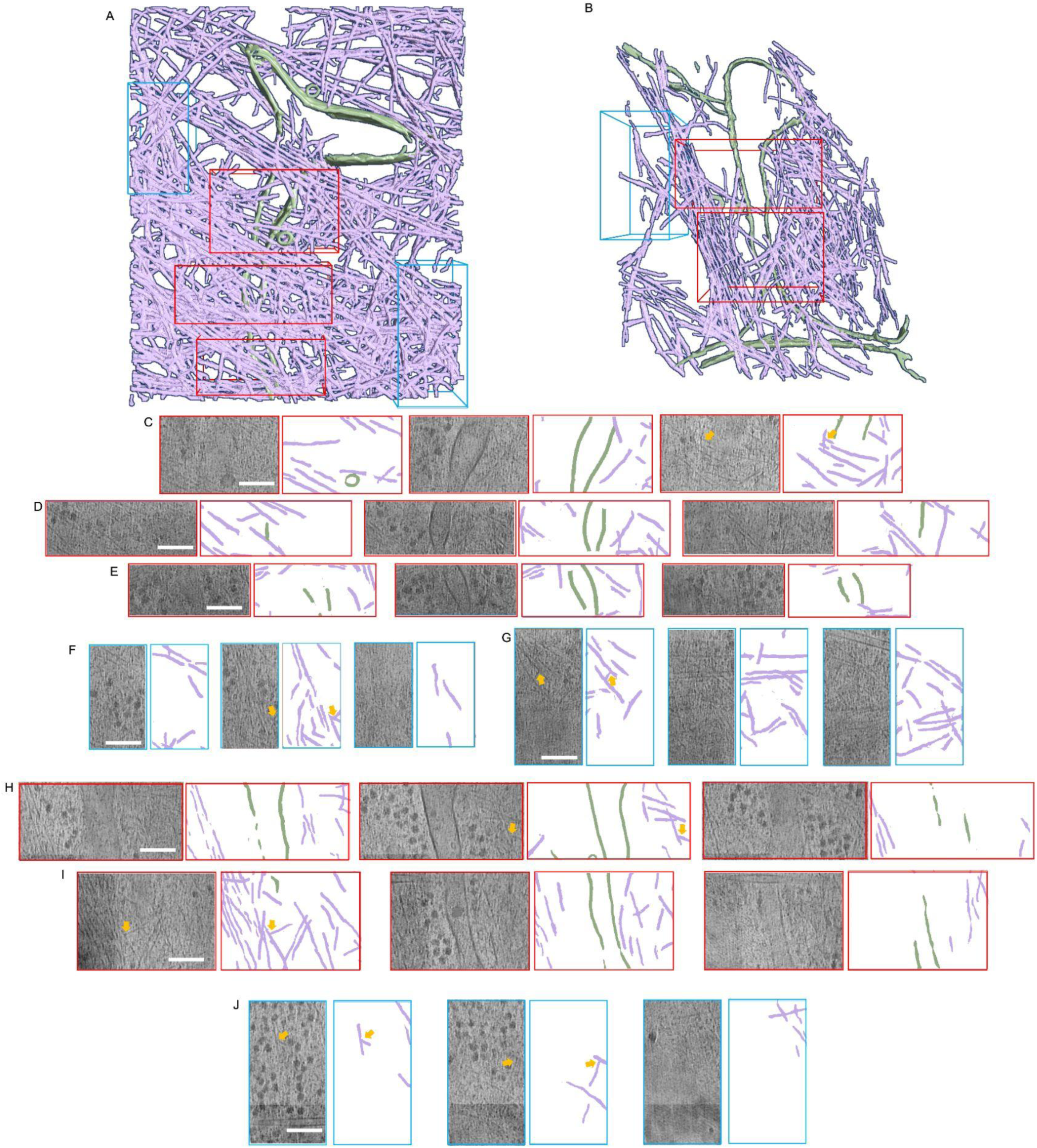
Quantitative analysis of cryo-electron tomograms. (**A**) 3D representation of cropped tomogram segmentation in fig S2E, sub-volumes proximal (red ROI) or distant (blue ROI) from tubule membrane used for counting branch number indicated with red boxes and blue boxes, respectively. (**B**) 3D representation of cropped tomogram segmentation in fig. S2F, sub-volumes proximal to and distant from tubule membrane used for counting branch number indicated with red boxes and blue boxes, respectively. (**C**) Slices and segmentations of the tomogram in A in red box1 at different z positions (from left to right: z=-35nm, 5nm and 34nm relative to the central slice) with actin filaments in purple, tubules in green, and potential actin branches indicated with yellow arrows. Scale bar: 100 nm. (**D**) Slices and segmentations of the tomogram in A in red box2 at different z positions (from left to right: z=-26nm, 3nm and 28nm relative to the central slice) with actin filaments in purple, tubules in green.Scale bar: 100 nm. (**E**) Slices and segmentations of the tomogram in A in red box3 at different z positions (from left to right: z=1 nm, 14 nm and 34nm relative to the central slice) with actin filaments in purple, Tubules in green.Scale bar: 100 nm. (**F**) Slices and segmentations of the tomogram in in A blue box1 at different z positions (from left to right: z=-27 nm, 3 nm and 22 nm relative to the central slice) with actin filaments in purple, tubules in green, and potential actin branches indicated with yellow arrows. Scale bar: 100 nm. **(G)** Slices and segmentations of the tomogram in A in blue box2 at different z positions (from left to right: z=-13nm, 10nm and 31nm relative to the central slice) with actin filaments in purple, tubules in green, and potential actin branches indicated with yellow arrows. Scale bar: 100 nm. **(H)** Slices and segmentations of the tomogram in B in red box1 at different z positions (from left to right: z=-15nm, 11nm and 34nm relative to the central slice) with actin filaments in purple, tubules in green, and potential actin branches indicated with yellow arrows. Scale bar: 100 nm. **(I)** Slices and segmentations of the tomogram in B in red box2 at different z positions (from left to right: z=-22 nm, 7 nm and 30 nm relative to the central slice) with actin filaments in purple, tubules in green, and potential actin branches indicated with yellow arrows. Scale bar: 100 nm. (**J**) Slices and segmentations of the tomogram in B in blue box at different z positions (from left to right: z=-14 nm, 16 nm and 39 nm relative to the central slice) with actin filaments in purple, tubules in green, and potential actin branches indicated with yellow arrows. Scale bar: 100 nm.

**Fig. S5.**
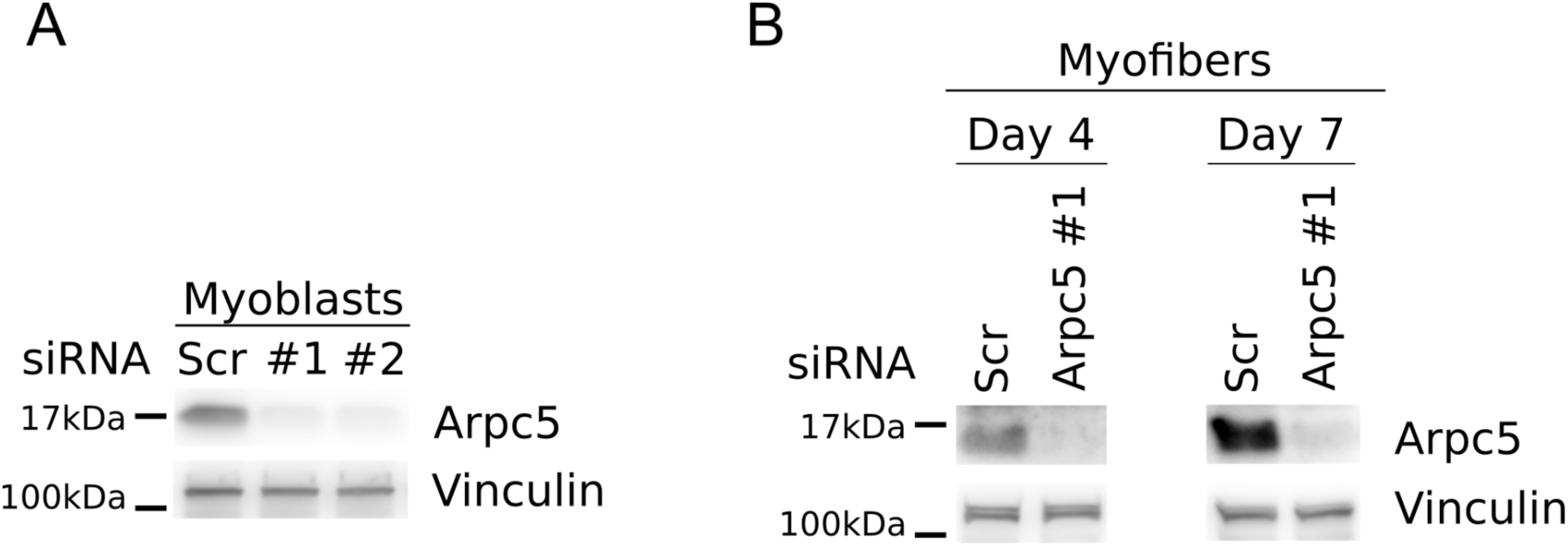
Western blots of Arpc5 depletion. (**A**) Western blots of GFP-Bin1 myoblasts knocked down for scramble (Scr), Arpc5 #1 (ID#s85623) or Arpc5#2 (ID#s85621). Membranes were blotted using antibodies against Arpc5 with vinculin as a loading control. (**B**) Western blots of myofibers (differentiation day 4 or 7), knocked down for scramble (Scr) or Arpc5 #1. Membranes were blotted using antibodies against Arpc5 and vinculin, used as a loading control.

## Movies

**Movie S1** – Timelapse movie of t-tubule dynamics in a myofiber isolated from the EDL muscle of an adult mouse. Isolated fiber was stained with CellMask to label t-tubules. Scale bar 5 μm.

**Movie S2** – Timelapse movie of t-tubule dynamics in a developing in vitro myofiber. Cells were stained with CellMask to label t-tubules. Scale bar 5 μm.

**Movie S3** – Timelapse movie of Triadin (green) and t-tubule (magenta) dynamics in a developing in vitro myofiber. Cells expressing Triadin-GFP stained with CellMask to label t-tubules. Scale bar 5 μm.

**Movie S4** – Timelapse movie of t-tubule dynamics in a developing myofiber before (left panel) or 5 min after (right panel) adding CK666. Cells were stained with CellMask to label t-tubules. Scale bar 5 μm.

**Movie S5** – Timelapse movie of tubule dynamics in GFP-Bin1 myoblasts before and after adding CK666. Scale bar 5 μm.

**Movie S6** – F-actin (LifeAct-mCherry, magenta) and Bin1 tubules (GFP-Bin1, green) in GFP-Bin1 myoblasts. Scale bar 5 μm.

**Movie S7** – F-actin (mScarlet-i-UtrCH, magenta) and Bin1 tubules (GFP-Bin1, green) in GFP-Bin1 myoblasts. Scale bar 5 μm.

**Movie S8** – Cryo-electron tomogram and segmentation model of actin cytoskeleton around t-tubule site. Actin cytoskeleton coloured in light purple. T-tubule membrane coloured in green.

**Movie S9** – Cryo-electron tomogram and segmentation model of actin cytoskeleton around t-tubule site. Actin cytoskeleton coloured in light purple. T-tubule membrane coloured in green.

**Movie S10** – Cryo-electron tomogram and segmentation model of actin cytoskeleton around t-tubule site. Actin cytoskeleton coloured in light purple. T-tubule membrane coloured in green.

**Movie S11** – Timelapse movie of tubule dynamics in GFP-Bin1 myoblasts before and after adding para-aminoblebbistatin (+Blebb). Scale Bar 5 μm.

**Movie S12** – Timelapse movie of t-tubule dynamics in a developing in vitro myofiber before (left panel) or 5 min after (right panel) adding para-aminoblebbistatin (+Blebb). Cells were stained with CellMask to label t-tubules. Scale bar 5 μm.

**Movie S13** - Timelapse movie of tubule dynamics in GFP-Bin1 myoblasts before and after adding glycerol. Scale Bar 5 μm.

**Movie S14** - Timelapse movie of t-tubule dynamics in a developing myofiber before (left panel) or after (right panel) adding glycerol. Cells were stained with CellMask to label t-tubules. Scale bar 5 μm.

**Movie S15** - Timelapse movie of tubule dynamics in GFP-Bin1 myoblasts before and after adding NSC668394 (+NSC). Scale Bar 5 μm.

**Movie S16** – Timelapse movie of t-tubule dynamics in a developing in vitro myofiber before (left panel) or 5 min after (right panel) adding NSC668394 (+NSC). Cells were stained with CellMask to label t-tubules. Scale bar 5 μm.

**Movie S17** – Timelapse movie of tubule dynamics in GFP-Bin1 myoblasts depleted for scramble control (left panel) or depleted for Arpc5 siRNA#1. Scale bar 5 μm.

**Movie S18** – Timelapse movie of t-tubule dynamics in a developing myofiber in scramble cells (top panel) or in cells depleted for Arpc5 (bottom panel). T-tubules were stained with CellMask. Scale bar 5 μm.

